# Insights into the domestication of avocado and potential genetic contributors to heterodichogamy

**DOI:** 10.1101/2022.03.30.486474

**Authors:** Edwin Solares, Abraham Morales-Cruz, Rosa Figueroa Balderas, Eric Focht, Vanessa E. T. M. Ashworth, Skylar Wyant, Andrea Minio, Dario Cantu, Mary Lu Arpaia, Brandon S. Gaut

## Abstract

- The domestication history of avocado (*Persea americana*) remains unclear, in part due to a lack of suitable genomic tools.
- We created a reference genome from the Gwen varietal, which is closely related to the economically dominant Hass varietal. We also compiled a database of 34 resequenced accessions that represented the three botanical races of *P. americana*.
- Our genome assembly had an N50 of 3.37 megabases, a BUSCO score of 91% and was scaffolded with a genetic map, producing 12 pseudo-chromosomes with 49,450 genes. We used the Gwen genome as a reference to investigate the population genomics of avocado. Our analyses were consistent with three separate domestication events; we estimated that the Mexican race diverged from the Lowland (formerly known as ‘West Indian’) and Guatemalan races >1 million years ago. We also identified putative targets of selective sweeps in domestication events; within the Guatemalan race, putative candidate genes were enriched for fruit development and ripening. We also investigated divergence between heterodichogamous flowering types.
- With the help of a new reference genome, we inferred the domestication history of avocado and identified genes that may contribute to heterodichogamy, including genes with functions in pollination and floral development.

## INTRODUCTION

Avocado (*Persea americana* Mill.) is a perennial, subtropical crop that is in ever-increasing demand. In the United States, for example, per capita avocado consumption has tripled over the last two decades. Demand in the U.S. is met partly by domestic production but principally by imports from Mexico and elsewhere. Mexico is the largest producer, where the crop is worth an estimated $2.5 billion per year (Rendón-Anaya *et al*., 2019), but other major producers include the Dominican Republic, Peru, Chile, Indonesia, Israel and Kenya (http://www.fao.org/faostat/en/#data/QC/visualize). Although the popularity of avocados is primarily a 20th century phenomenon (Schaffer *et al*., 2013), they have quickly grown to be a global commodity.

Remarkably, avocado cultivation is dominated by a single variety (Hass) that represents ~90% of cultivation world-wide (Rendón-Anaya *et al*., 2019). All Hass trees are derived clonally from a tree patented in 1935. Despite the shockingly narrow genetic base of agricultural production, avocado *sensu lato* is quite genetically diverse. Some of this diversity stems from the fact that there are three domesticated botanical races: *P. americana* var. *americana* Miller ([which we will call the ‘Lowland’ race in recognition that the previously accepted name of West Indian is inaccurate (Me & Arzate-Fernández, 2010)], var. *drymifolia* (Schltdl. & Cham.) S.F. Blake (the Mexican race), and var. *guatemalensis* (L.O. Williams) Scora (the Guatemalan race) (Bergh & Ellstrand, 1986). The strikingly different fruit morphologies among the races suggest that they may have been domesticated separately, a conjecture supported by genetic data (Furnier *et al*., 1990; Ashworth & Clegg, 2003; Rendón-Anaya *et al*., 2019). One practical consequence of this history is that each race likely contains separate alleles and/or genes of interest for crop improvement, due to their different domestication histories. Another consequence is that hybridization between races can produce unique allelic combinations, potentially leading to agronomically useful hybrid offspring. Hass may be, in fact, one example; although its precise breeding history is not known, genetic evidence has suggested that it is a hybrid between Guatemalan and Mexican races (Chen *et al*., 2009; Schaffer *et al*., 2013; Rendón-Anaya *et al*., 2019).

The high demand for, and economic importance of, avocados motivates breeding efforts, but breeding remains challenging for at least three reasons. First, avocado is a large tree that matures slowly [5 to 8 years before production (Lahav & Lavi, 2009)] requiring substantial space, water and finances (Ashworth *et al*., 2019). A second major obstacle is the reproductive system. A single tree typically produces more than one million flowers, of which only 0.1% or fewer yield mature fruit (Davenport, 1986; Davis *et al*., 1998), making controlled pollinations difficult (Chen *et al*., 2007). Finally, avocado is predominantly out-crossing, due to heterodichogamy. There are two flowering types in the avocado system: A and B. Type A trees are female (receptive to pollen) in the morning of the first day and shed pollen as males in the afternoon of the following day. In contrast, type B trees are female in the afternoon of the first day and male in the morning of the next day. Although heterodichogamy is likely caused by a simple underlying genetic mechanism (Renner, 2001), the system is complicated by the fact that there is some leakiness that depends on environmental cues (Davenport, 1986). As a result of these complications, avocado breeding has historically relied on open-pollinated and inter-racial hybridization, to the extent that most individual varieties lack accurate breeding records (Davis *et al*., 1998; Scora *et al*., 2002; Ashworth & Clegg, 2003).

These complications argue that genomics and molecular breeding are central for the continued improvement of avocado. Molecular markers for flowering types may be particularly useful, because type B avocados are crucial for pollination but typically less productive than type A varieties (Davenport, 1986). Recently, Rendón-Anaya et al. (2019) made an important contribution toward molecular breeding by producing draft genomes of Hass and a wild Mexican accession (*P. americana* ssp. *drymifolia*). They anchored the Hass assembly to a genetic map and ultimately produced a reference genome with 512 scaffolds and a genome size of 419 Megabases (Mb), which is less-than-half the expected 1C genome size of 896Mb (Arumuganathan & Earle, 1991). Using this reference, they explored the hybrid history of Hass and aspects of the evolutionary genomics of avocado. Another Hass genome was published in 2021 (Sharma *et al*., 2021); this genome was 788Mb, which was nearly 88% of the expected genome size, but it was not anchored into scaffolds. Despite the availability of these genomes, several important features of the evolutionary genomics of avocado remain unexplored, including further unraveling its domestication history, using evolutionary genomic tools to identify chromosomal regions of potential agronomic interest, and focusing on genomic diversity in the context of interesting traits, like A vs. B flowering types.

Here we report the genome of the Gwen variety and use that genome as a reference for evolutionary analyses. Gwen is a grandchild of Hass with similar flavor characteristics (Witney & Martin, 1995) but with higher yields and better fruit storage on the tree (Bergh & Whitsell, 1982). Accordingly, Gwen has been the subject of intensive breeding efforts for three decades.

We also generated a whole-genome resequencing of 34 avocado varieties to help focus on four sets of issues. First, what does the Gwen genome tell us about patterns of genic hemizygosity within an avocado accession? Genomic analysis of grapevine (*Vitis vinifera*),another perennial clonally propagated crop, revealed that as many as one in seven genes are hemizygous, perhaps due to structural mutations that have accrued during clonal propagation (Zhou *et al*., 2019). Is avocado similar? Second, we use the Gwen genome as a reference to explore genetic diversity within avocado, specifically to assess relationships among the three races and to assess the hybrid origin of well-known cultivars. This last question builds on several previous investigations of genetic diversity (Ashworth & Clegg, 2003; Chen *et al*., 2008, 2009) but extends the work to a genomic scale. Third, we investigate features of avocado domestication, including demographic history, selective sweeps and chromosomal regions of high genetic differentiation. Do the three racial groups share regions of selective sweeps, which may indicate parallel selection on genic regions associated with specific traits? Finally, we investigate genetic diversity between the A and B flowering types, with the goal of identifying genomic regions that may contribute to heterodichogamy.

## MATERIALS AND METHODS

### Reference Sequencing and Genome Assembly

Gwen (US Patent: USPP5298P 1983) is a grandchild of Hass and has been central to the University of California, Riverside breeding program. We sampled young leaf materials from a Gwen clone and isolated high molecular weight genomic DNA (gDNA) using the method of Chin *et al*., (2016). We included a containing buffer (100 mM Tris-HCl pH 8.0, 0.35 M Sorbitol, 5 mM EDTA pH 8.0, 1% (v/v) PVP 40, 2% (v/v) of 2-mercaptoethanol) prior to cell lysis to avoid co-precipitation of polysaccharides and phenolics with DNA. DNA quality was assessed with a Nanodrop 2000 spectrophotometer (Thermo Scientific, IL, USA), by a Qubit 2.0 Fluorometer with the DNA High Sensitivity kit (Life Technologies, CA, USA), and by pulsed-field gel electrophoresis. From this, 20 μg of high molecular weight genomic DNA was needle sheared and used as input into the SMRTbell library preparation, using the SMRTbell Express Template Preparation V2 kit (Pacific Biosiences, CA, USA). Libraries were size selected to 22-80Kb with the BluePippin (Sage Science, MA, USA) and sequenced on the Pacific Biosciences Sequel using P6-C4 chemistry (DNA Technology Core Facility, University of California, Davis).

We assembled SMRT reads with Canu version 2.1, using default settings and including all reads. Once assembled, polishing was performed with two passes of PacBio GenomicConsensus v2.33, followed by two passes with Pilon v1.23 (Walker *et al*., 2014) using default parameters and 19x coverage of Gwen short read Illumina sequencing data. HapSolo v0.1 (Solares *et al*., 2021), which identifies and removes alternative haplotypes, was then run on the assembly with default parameters and 50,000 iterations, producing the Canu+Hapsolo (C+H) assembly. Scaffolding was based on a Gwen x Fuerte genetic map (Ashworth *et al*., 2019) by aligning the C+H assembly using NCBI BLAST v2.2.31+ (Altschul *et al*., 1990). The alignments were then ordered based on linkage group ID and cM distance, using only the alignments with the highest percent identity and evalue scores to identify contig order. When the orientation of a contig could not be determined, it was placed in the ‘+’ (or positive) direction and marked with an asterisk in the scaffolding annotation file (10.5281/zenodo.6392169). Where necessary, contigs were bridged using N’s.

### Gene and Repeat Annotation

Repeat annotation was based on RepeatModeler v2.0.1 in conjunction with RepeatMasker v4.1.1. RepeatModeler was run to generate a repeat database for avocado, since a database for closely related species was not available. The repeat database was built using the option BuildDatabase on the Gwen H+C assembly and then used in RepeatModeler with the LTRStruct option. RepeatMasker was run using default parameters. The Gwen genome was subsequently soft masked for repeats using Bedtools v2.29.2 using the maskfasta run option.

For gene annotation, we mapped RNASeq reads from previous studies (Ibarra-Laclette *et al*., 2015; Xoca-Orozco *et al*., 2017; Barbier *et al*., 2019) onto the repeat masked genome using HiSat2 version 2.2.1. The resultant BAM files were merged and indexed using Samtools v1.10 (Li *et al*., 2009; Barnett *et al*., 2011) and fed to the BRAKER v2.1.6 (Hoff *et al*., 2019) /Augustus (Stanke *et al*., 2006) v3.4.0 pipeline. BRAKER was used in default mode for RNASeq data, with an additional option for softmasked reference assemblies (--softmasking). Finally, we excluded (as possible pseudogenes) any genes with exons that overlapped annotated repeats and filtered for specific criterion related to gene integrity (Minio *et al*., 2021).

Functional annotations and Gene Ontology analyses were performed with Blast2GO v6.01 (Conesa & Götz, 2008). Genes were extracted from assemblies, based on the gene annotation gff file and then mapped to the NCBI non-redundant protein database, SwissProt and uniref90 databases using NCBI’s blastx, blastx-fast option. A Protein family search was also performed using an InterPro scan. These results were then merged into a single annotation for GO mapping and Enrichment, which was performed using Blast2GO based on default options.

Genic hemizygosity was calculated following Zhou et al. (2019) by: *i*) remapping raw SMRT reads to the 12 pseudo chromosomes using NGMLR v0.2.7, *ii*) inferring structural variants by feeding the alignment file to SNIFFLES (Sedlazeck *et al*., 2018), requiring a read-coverage of at least four to substantiate an SV, and *iii*) and quantitating insertion deletion events that overlapped with 20% of the coding region.

### Diversity Samples, Sequencing and SNP calling

We collected leaf tissue for a total of 20 *P. americana* accessions and three outgroups (**Table 1**), all of which were sampled from the South Coast Research and Extension Center in Irvine, CA. For each sample, genomic DNA was extracted from leaf samples with the Qiagen DNeasy plant kit. Paired-end sequencing libraries were constructed with an insert size of 300 bp according to the Nextera Flex (Illumina, Inc) library preparation protocol. Libraries were sequenced on the Illumina NovaSeq with cycles to target 25X coverage. We also used Illumina reads for ten previously sequenced *P. americana* accessions and one putative outgroup (Rendón-Anaya *et al*., 2019). Short reads were mapped to the H+C and scaffolded assemblies using BWA version 0.7.17 (Li & Durbin, 2010) and realigned and recalibrated using GATK v3.7 (McKenna *et al*., 2010). Alignment filtering was done using BCFTools v1.10.2 (Danecek *et al*., 2021) using parameters *-S LowQual -e* ‘*%QUAL*<20 || *DP*>32′.

**Table 1:**
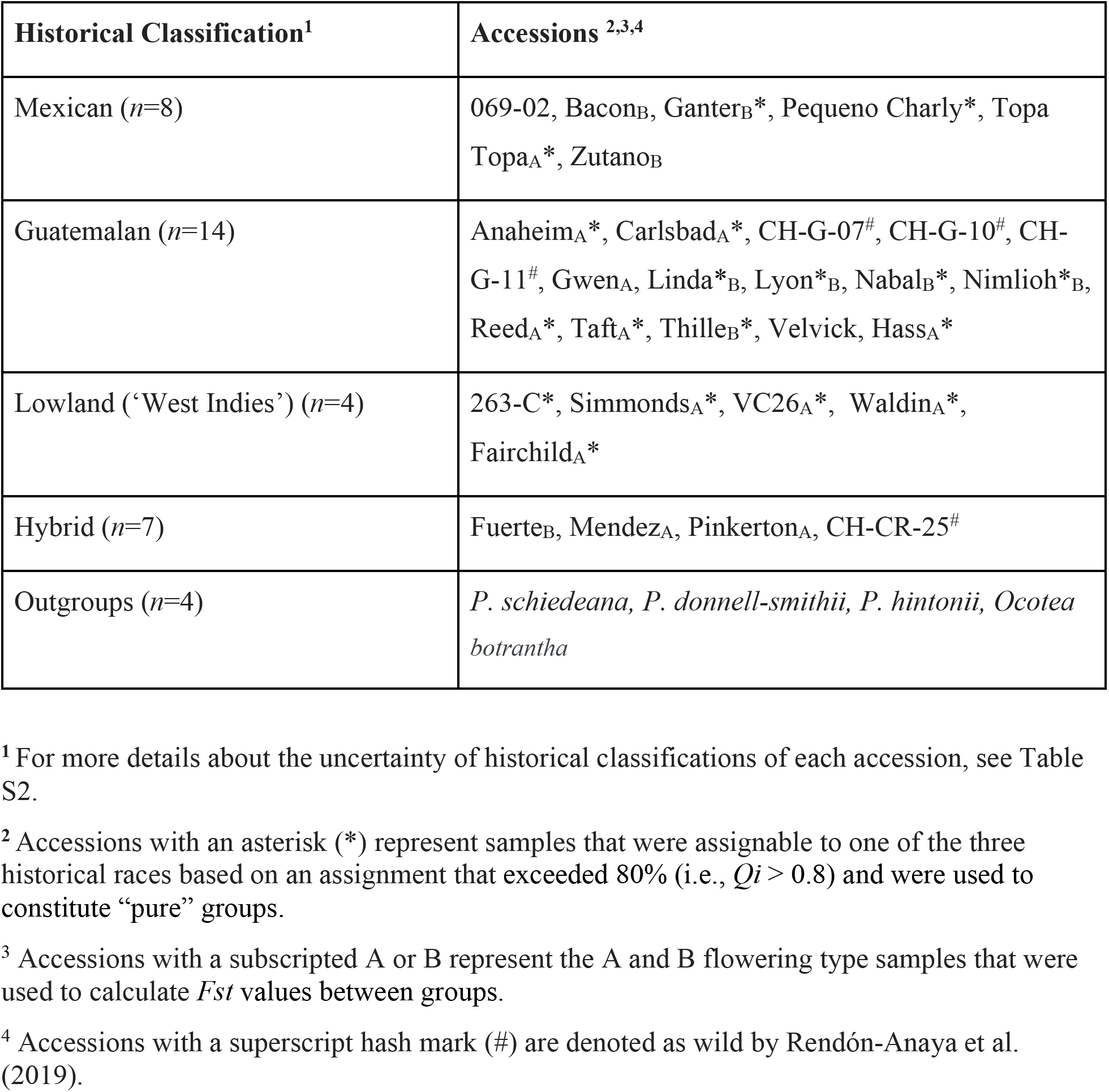
A list of accessions used in diversity studies. Additional details are provided in Table S2.

### Phylogeny and Population Structure

We performed PCA using VCFTools v0.1.17 git commit 954e607 (Danecek *et al*., 2011), and PLINK v1.9 (Purcell *et al*., 2007). All samples had < 50% missing data for *P. americana* and <75% for outgroup taxa. We created a maximum likelihood phylogeny with a reduced number of sites from SNPhylo (Lee *et al*., 2014) using IQ-TREE v1.6.12 (Nguyen *et al*., 2015) and the “PMB+F+G4” substitution model, as chosen by Model Finder (Kalyaanamoorthy *et al*., 2017). We employed the ultrafast bootstrap with 1000 replicates to obtain support values. Admixture plots and analyses were performed with NGSadmix from the ANGSD package version 0.930 (build Jan 6, 2020 13:30:06). CLUMPAK (Kopelman *et al*., 2015) was used for identified the best number across K=1 to 10 groups, each with 10 replicates.

### Population Genetic Analyses

To infer demographic histories, we used MSMC2 v2.1.1 (Mallick *et al*., 2016) with phased SNPs from the entire scaffolded assembly, which included the 12 pseudo-chromosomes and unplaced contigs. For each genetic group, we applied MSMC2 to the three accessions with the highest average sequencing coverage per genetic group (Guatemalan: Lyon, Carlsbad, Nimlioh; Lowland: Fairchild, Waldin, VC26; Mexican: Ganter, PequeñoCharly, TopaTopa). We created a mappability mask and a coverage mask (**Methods S1**). Phase-informative SNPs (PIRs) were extracted for each sample as described in (Delaneau *et al*., 2013), which identifies reads that span at least 2 heterozygous SNPs; we then used the PIRs and the VCF files for each genetic group by chromosome as input to shapeit2 v2.r904 (Delaneau *et al*., 2013) for phasing. Finally, we used the “generate_multihetsep.py” script from the MSMC tools (https://github.com/stschiff/msmc-tools) to create MSMC2 input, taking into account mappability and coverage masks. To calculate the relative cross coalescence rate (rCCR) across groups we ran MSMC2 with all possible haplotype combinations between groups. To calculate split times, we used “plot_msmc.py” (Schiffels & Wang, 2020) to estimate time at rCCR=0.5, based on a rate of 5.4e-9 mutations per site per year (Liang *et al*., 2019) and a generation time of 7 years.

To infer selective sweeps, we applied SweeD (Pavlidis *et al*., 2013) with default parameters on the scaffolded assembly, including unplaced contigs. To perform the analyses, we split the genome into non-overlapping 10 kb windows created by bedtools v2.27.1 (Quinlan, 2014), ignoring gaps, and focused on windows within the top 1% of the Composite Likelihood Ratio (CLR) statistic. SweeD identifies the location of putative sweeps; to identify genes encompassed in that sweep, we included genes 5kb on either side of the location. Once genes were identified, we performed two types of analyses: GO enrichment, as described above, and the statistical significance of the number of shared sweep genes between races. To evaluate significance, we permuted labels on genes (either sweep or non-sweep) within each race, recalculated the number of sweep genes in common between races, and repeated the permutation 10,000 times. We used PLINK (Purcell *et al*., 2007) on 20kb windows to calculate mean *Fst* for each window between samples, focusing on the top 1% windows.

## RESULTS

### Gwen Genome Assembly and Characterization

#### Gwen genome assembly

We generated 81.2 giga bases, equivalent to roughly 90x coverage, based on the expected 1C genome size of 896Mb (Arumuganathan & Earle, 1991). We assembled PacBio SMRT reads using Canu (v 2.1), producing a genome of 1,456Mb with 5,122 contigs and then applied HapSolo (Solares *et al*., 2021) to remove putative secondary contigs (or haplotigs). The Canu+HapSolo (C+H) genome resulted in a primary assembly of 1,032 Mb, a longest contig of 17Mb, a BUSCO score of 91% and an N50 of 3.37Mb (**Table S1**). One useful measure of an assembly is the percentage of the assembly that is encompassed in the largest *x* contigs, where *x* represents the number of chromosomes (which is 12 for 1C *P. americana*). For the C+H assembly, this percentage was 17% - i.e., the 12 largest contigs represented 17% of the genome.

To improve contiguity, we anchored and scaffolded the C+H assembly using a published genetic map, based on a cross between Gwen and Fuerte (Ashworth *et al*., 2019). This exercise resulted in 12 scaffolds that were assigned to 12 linkage groups, along with unplaced contigs. Scaffold N50 improved ~18-fold (to 61.9Mb) over the 3.37Mb contig N50, and the 12 largest scaffolds (pseudo-chromosomes) represented 78% of the expected genome size of 896Mb (**Table S1**). Although each chromosome could be identified among the 12 largest scaffolds, we interpret the smaller-than-expected genome size to imply that the density of the genetic map limited resolution. The first Hass primary assembly also decreased substantially in size when it was anchored to a genetic map (Rendón-Anaya *et al*., 2019) - e.g., only 47% of the Hass assembly could be anchored, resulting in a scaffolded genome of 421.7Mb, while the second lacked any scaffolding (Sharma *et al*., 2021). Hence our scaffolded genome is nearly double the size of the previous scaffolded avocado reference.

We compared our new Gwen genome to the two existing Hass assemblies and to the *P. drymifolia* assemblies. The Gwen assembly was superior for N50, longest contigs (or scaffolds), and the proportion of the genome captured in the 12 largest fragments (**Table S1**). We also plotted the cumulative density function of contigs and scaffolds for the four genomes (**Figure 1A**), providing a virtual comparison among the genomes. While 78% of the Gwen assembly was captured in the largest 12 scaffolds (or contigs), this value was 4.2% and 12.8% for the two Hass assemblies and 2.7% for the *drymifolia* assembly (Rendón-Anaya *et al*., 2019; Sharma *et al*., 2021).

**Figure 1:**
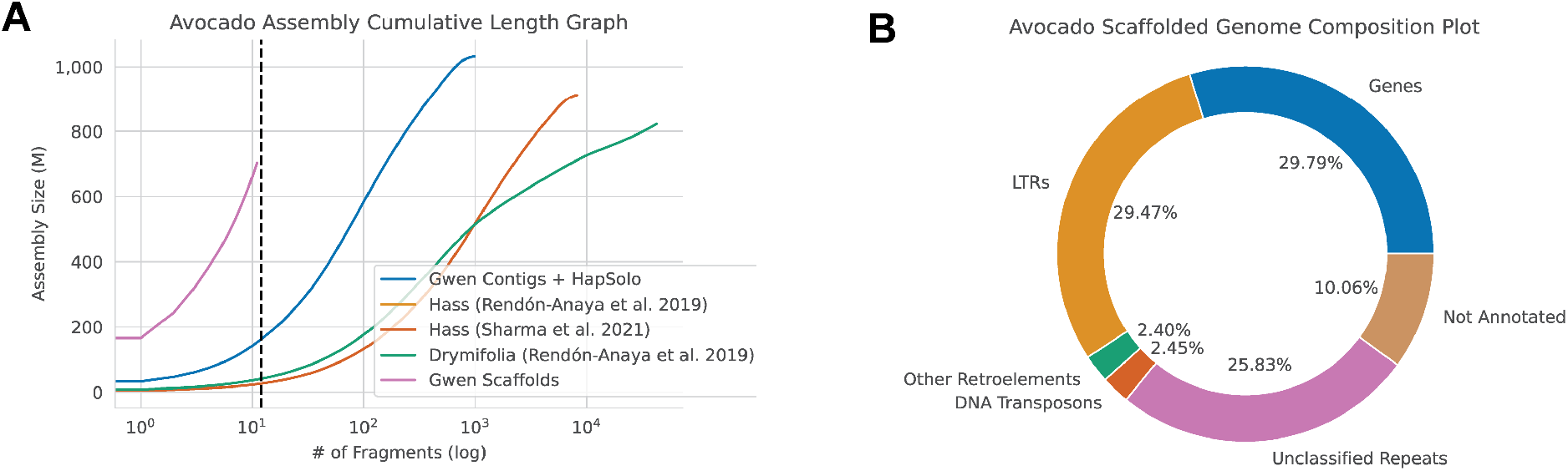
(**a**) The cumulative sum assembly graph size (cdf) shows the size of the assemblies with the next largest consecutive contig (or scaffold) being added to the sum along the *x*-axis. It illustrates cdf for the Hass and *drymifolia* assemblied from Rendon-Anaya et al. (2019) and the Hass HiFi assembly from Sharma et al. (20210), and the two Gwen assemblies (C+H and scaffolded). (**b**) The annotation results for the Gwen C+H assembly, showing the percentage of the genome attributable to genes and different types of transposable elements, including DNA transposons, unclassified retroelements and Long Terminal Repeat (LTR) retroelements.

#### Genome Annotation

We annotated the C+H assembly and scaffolded assemblies by first identifying the repetitive content (see Methods). We estimated that ~61% of the Gwen genome consisted of repetitive elements (**Figure 1B**), of which 52% were Long Terminal Repeat (LTR) retroelements (with more than half of these being *Gypsy* elements) and another 40% were unknown repeats (**Figure 1B**). We masked repeats before annotating genes (see Methods), inferring 36,993 genes on the 12 pseudo-chromosomes and 12,457 in unplaced contigs. We compared these numbers to the Hass genome annotation (Rendón-Anaya *et al*., 2019), which reported 33,378 genes. We filtered this set to remove potential duplicates, yielding 25,211 genes. Of the 25,211, 94% were present in our scaffolded assembly; thus, our annotation corroborates most previous genic inferences but also annotates ~2-fold more genes.

#### Genic Hemizygosity

Recent studies have suggested that diploid genomes may be replete with genic hemizygosity. We assessed genic hemizygosity by remapping raw PacBio reads to the 12 pseudo-chromosomes of the scaffolded assembly, by inferring structural variants and then by filtering for variants that overlapped annotated genes (see Methods). With this approach, we estimated that 3.8% of genes were hemizygous in the Gwen genome. This estimate serves as a benchmark that can be used to compare to other diploid systems (see Discussion).

### Analysis of *P. americana* Genetic Structure

#### Classification of races and hybrid varieties

Given the Gwen reference genome, we performed a preliminary study of genetic variation and evolutionary genomics across avocado. To do so, we amassed a whole-genome resequencing dataset of 30 *P. americana* accessions and four putative outgroups with at least 14x coverage (**Table 1**). The sample represented the three botanical races of avocado, based on historical designations, historically important cultivars (Chen *et al*., 2009), and both A and B flowering types (**Table 1 & S2**). Given the resequencing data, we mapped reads to the scaffolded assembly (including unplaced contigs) and identified 23,938,386 biallelic SNPs within our entire *P. americana* sample and 32,235,263 biallelic SNPs with outgroups.

SNPs were then subjected to three types of clustering analysis – principal components (PCA), admixture mapping and phylogenetic inference -- to define groups of accessions based on whole genome data. We first applied the PCA to the *P. americana* data without outgroups. The results verified many of the expected groupings, including clusters that the represented previously identified Mexican, Guatemalan and Lowland races (**Figure 2A**). There were, however, several accessions that were located between groups on the PCA - e.g., Velvick - that suggest a hybrid origin (**Table S2**). One surprise came from the location of Hass and Gwen on the PCA. Hass clustered more closely to Guatemalan accessions than was expected of an accession that has been previously defined as >50% Mexican (Rendón-Anaya *et al*., 2019) and 58% Guatemalan by Chen *et al* (2009). Gwen is the grandchild of Hass and also clustered near Guatemalan accessions, suggesting again that it is primarily from race *guatemalensis*. Important varieties like Bacon and Zutano have been inconsistently inferred or assigned as hybrids (Chen *et al*., 2009) but appear to have had a hybrid origin based on whole genome analyses (**Figure 2A & Table S2**).

**Figure 2:**
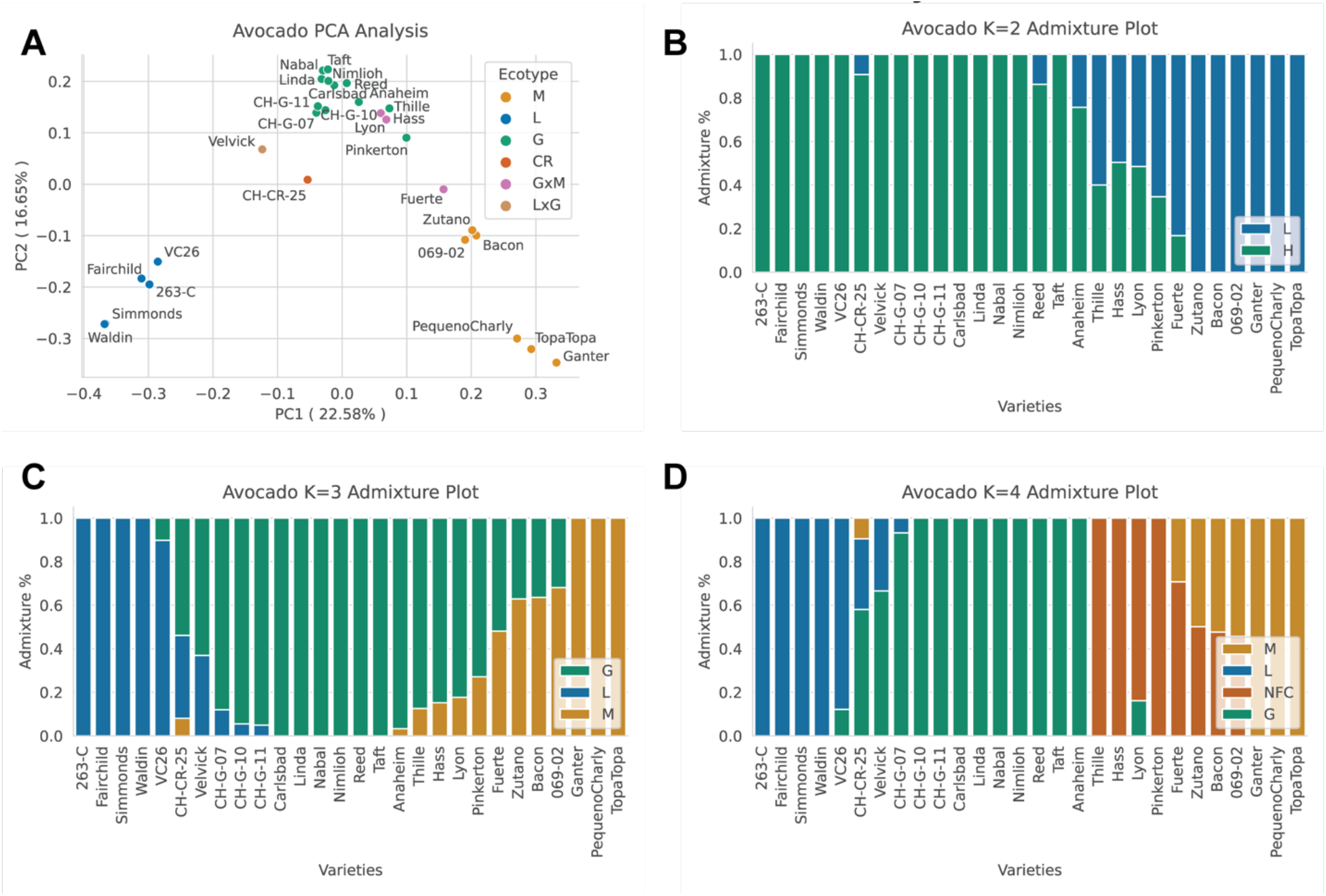
(**a**) A PCA analysis of SNP diversity, based on 29 accessions of avocado (without including the Gwen reference accession). Each dot represents an individual, and the color of each dot represents an historical classification of that individual, including Guatemalan x Mexican (GxM) hybrids and Lowland x Guatemalan (LxG) hybrids, and CH-CR-25, the putatively wild Costa Rican (CR) variant. (**b,c,d**) Admixture plots were generated using SNPs found in all of the contigs in the Gwen scaffolded assembly. Plot (**b**) shows the inferred groups with K=2 groups; plot (**c**) represents K=3, which is consistent with the three historical races; and (**d**) represents K=4 groups, which was inferred to be the most likely number of clusters. The K=4 groups include the three historical races and an emergent group showing accessions closely related to Hass.

For completeness, we also applied PCA to a dataset with the four outgroup taxa (**Figure S1**). The genetic placement of three species (*P. donnell-smithii* Mez*, P. hintonii* C.K. Allen*, O. botrantha* Rohwer) was consistent with their expected outgroup status, but *P. schiedeana* Nees often clustered within the avocado ingroup. These results are consistent with suggestions that *P. schiedeana* hybridizes with avocado (Ashworth & Clegg, 2003), indicating it should not be used as an outgroup for evolutionary analyses.

We also investigated relationships among accessions using admixture mapping. The most highly supported analysis contained k=4 groups: the three previously-recognized races (Lowland, Mexican, Guatemalan) and a series of cultivars related to Gwen and Hass (**Figure S2**). This last group included Mendez, a somatic mutation of Hass (Illsley-Granich *et al*., 2011; Schaffer *et al*., 2013), and other close relatives. We suspect that this last group was defined by over-sampling varieties with first- or second-degree relationships to Hass (including Hass, Gwen, Thille and Mendez). Indeed, removal of all but one of the ‘Hass group’ yielded K = 3 as the most supported number of groups, with the three groups representing the previously recognized races (**Figure 2C**). These admixture maps reinforced the hybrid history of some varieties, including Velvick, Fuerte, Bacon, Zutano and also an accession (CH-CR-25) that was previously thought to represent a new racial ecotype - var. *costaricensis* (Ben-Ya’acov *et al*., 2003; Rendón-Anaya *et al*., 2019).

Finally, we created a consensus phylogenetic tree to investigate relationships among races. We based this tree both on non-hybrid accessions that had high support (*Qi* > 80%) for inclusion in a single genetic group at K=3 (Vigouroux *et al*., 2008; Castillo *et al*., 2010) and also on a reduced number of SNPs to limit linkage disequilibrium (see Methods). The phylogeny had median bootstrap support of 88.5% for all nodes and strong support (> 76%) for nodes that separated the botanical races (**Fig. 3A**). Moreover, the accessions from each race formed well-supported monophyletic clades, justifying treating each named race as historically separate. An interesting feature of the rooted phylogeny is that the Mexican group is basal within the avocado ingroup, suggesting an early split from the Lowland and Guatemalan lineages and a separate domestication of Mexican avocado. Another interesting feature is the placement of putatively wild accessions, CH-G-07 and CH-G-10. These accessions intercalate the Lowland and Guatemalan clades, reinforcing the idea that the two races were domesticated separately (Furnier *et al*., 1990; Ashworth & Clegg, 2003; Rendón-Anaya *et al*., 2019).

**Figure 3:**
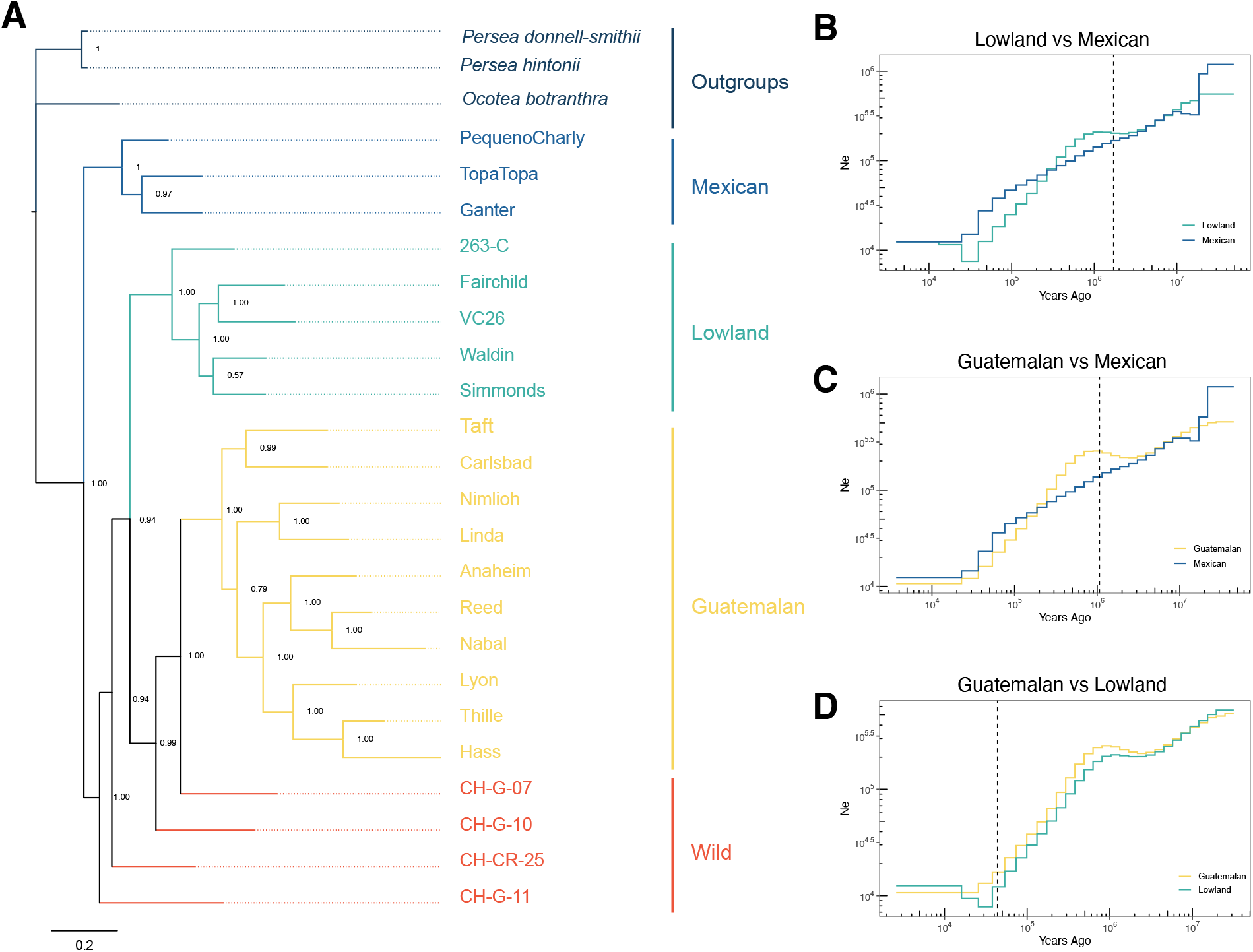
(**a**) Genome-wide phylogeny of *P. americana* ingroup with three outgroups. Accessions were chosen if they had 80% of higher assignments to a race, including putatively wild accessions (CH-G-07, CH-G-10, CH-G-11 and CH-CR-25). The numbers on the nodes represent bootstrap proportions. (**b**) Demographic inference based on Mexican and Lowland samples. The x-axis represents time from present to past, and the y-axis is inferred effective population size. The vertical dashed lines represented the estimated divergence time between the two races. (**c**) Like b, except for the Mexican and Guatemalan samples. (**d**) Like b, except for the Lowland and Guatemalan samples.

### Population Diversity Analyses

Our sampling was designed to include representative samples from each botanical race. However, analyses of genetic structure identified hybrids that were unlikely to be helpful for inferring historical dynamics of specific races. Accordingly, we focused population genetic analyses on samples with *Qi* > 80% (**Figure 3A**). This resulted in different sample sizes for the three races, with the Mexican sample particularly small (*n*=3) (**Table 1**). These sample sizes likely limit inferential power for some analyses, but we nonetheless retained > 10 million SNPs within each sample, with nucleotide diversity (π) similar among races, at ~0.0035 per nucleotide site (**Table S3**).

#### Demographic history

We applied the Sequentially Markovian Coalescent to infer two distinct aspects of the history of botanical races. The first was to infer whether any experienced a domestication bottleneck or other dramatic demographic event, recognizing that perennial crops often lack a signature of such events (Gaut *et al*., 2018). The results indicated a consistent reduction of *Ne* over time but without any evidence for a particularly notable bottleneck or rapid post-bottleneck expansion (**Figure 3B**). The second was to estimate the timing of the split of botanical races; the divergence times complemented the phylogeny by indicating an early split (~1.3 million years ago) of the Mexican group and with the Lowland and Guatemalan races diverging more recently, at ~44,000 years ago. These split-times predate expected domestication times and thus likely reflect divergence times between ancestral wild progenitor populations (**Figure 3B**).

#### Selective Sweep Mapping

For the three sets of samples representing potentially distinct domestication events, we investigated two additional features of their evolutionary genomics. The first was sweep mapping within each botanical race and the second was divergence between groups, as measured by *Fst*. We performed these analyses to assess whether similar (or entirely different) sets of genes bear marks of selection across races, to investigate whether highly differentiated chromosomal regions between races overlapped with selected genes, and to generate a list of candidate selected genes with potential functions.

Sweep mapping relied on the CLR statistic and focused on 10kb non-overlapping genomic windows of the scaffolded assembly. Using an empirical cut-off of 1%, we identified 1300 windows from each race, for which 638, 436 and 92 had genes in the Mexican, Lowland and Guatemalan samples, respectively (**Table S4**). Given these genes, we first hypothesized that separate domestication events may have targeted particularly important sets of genes in parallel. Visually, based on smoothed curves of the CLR statistic, this did not appear to be the case (**Figure 4**), because putative sweep regions differed markedly among samples. However, we also asked the question more formally by calculating the number of shared sweep genes between pairs of races and then permuting labels (CLR, non-CLR) to test significance. We found no enrichment for shared CLR genes between the Guatemalan sample and either the Mexican (p=0.3692; 2 shared genes; **Table S5**) or the Lowland samples (p = 1.00; 0 shared genes). However, the Lowland and Mexican groups had 18 CLR genes in common, a number significantly higher than random expectation (p < 0.0001) (**Table S6**).

**Figure 4:**
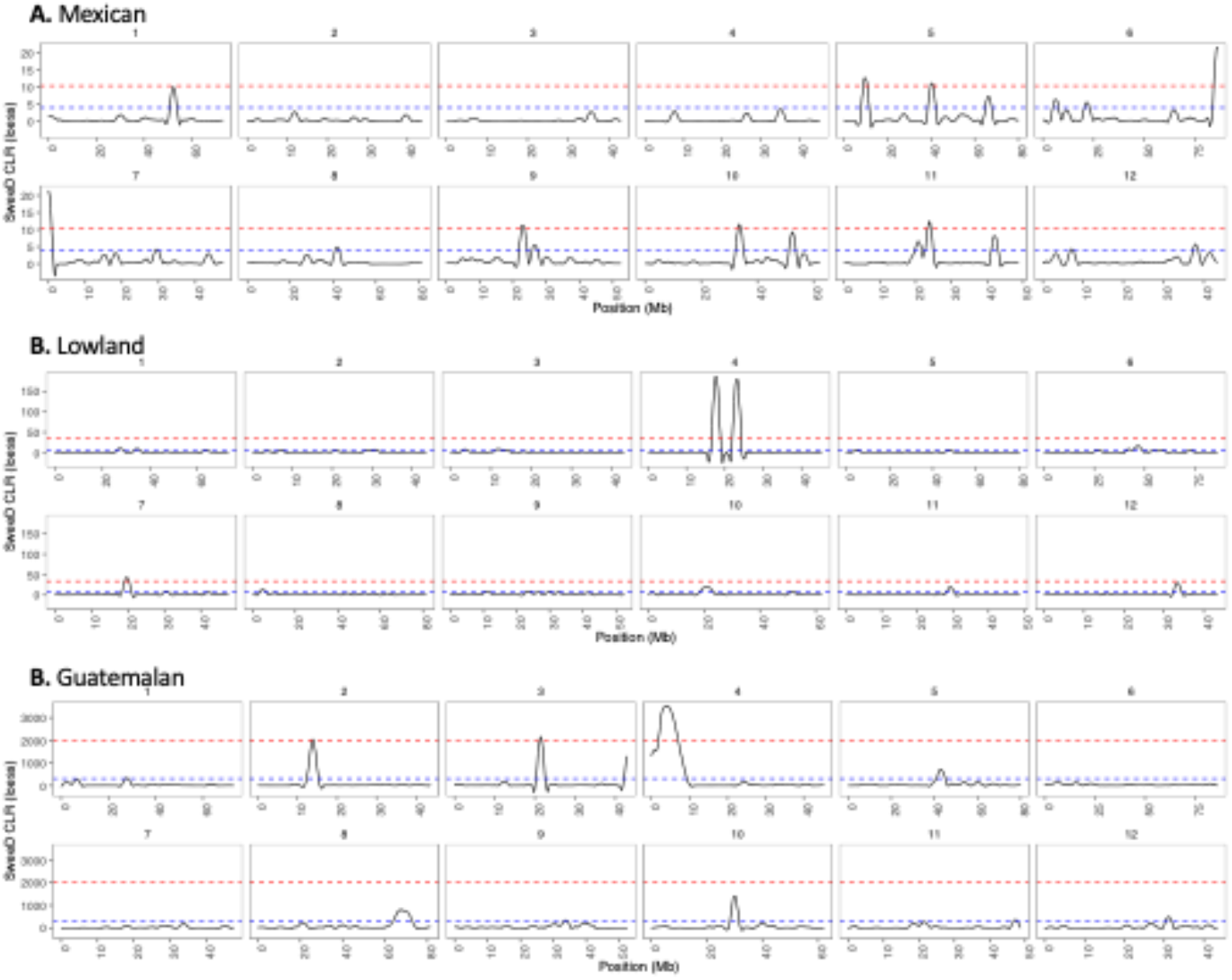
Loess smoothed composite likelihood values reflecting potential regions in samples from each of three botanical races: (**a**) Mexican, (**b**) Lowland and (**c**) Guatemalan. Each panel represents the 12 scaffolded chromosomes, with chromosome number and location (in Mb) provided above and below each individual graph. In each graph, the dashed lines represent 1% and 5% cutoff values for significance.

We evaluated sets of selected genes using GO enrichment. In the Guatemalan race, for example, the putative swept genes were enriched for functions related to fruit ripening, fruit development, anatomical structure maturation and other functions (**Figure S3**). Since these functional enrichments reflect properties potentially associated with domestication, we consider this genic set to be noteworthy. In contrast, we did not identify enrichment of gene function related to fruit maturation or suspected domestication traits for the Mexican and Lowland samples (**Figures S4 & S5)**, perhaps reflecting statistical uncertainty due to their small sample sizes. We did, however, identify gene enrichments related to functions like stress response, terpene production and metabolic processes. The set of shared 18 genes was also particularly interesting, due to the potential for parallel selection during separate domestication events. A subset of the genes had positive hits to functional annotation databases (**Table S6**). However, with the possible exception of two genes that function in sugar transport, none have functions related to obvious domestication traits like fruit size or development.

#### Divergence Mapping

We next examined divergence between races using *Fst* based on 20kb non-overlapping windows along scaffolded chromosomes, focusing on regions that include the top 1% of *Fst* scores (**Figure S6**). For example, for the pairwise comparison between Mexican and Lowland samples, we identified peaks containing 396 genes, with similar numbers for the other pairwise comparisons (Mexican-Guatemalan 387 genes; Lowland-Guatemalan 384 genes) (**Table S7**). In contrast to the potential for parallel domestication pressures on some genes, we expected *Fst* results to be enriched for genes that contribute to agronomic differences between the two races. Several genes in the top 1% were related to disease resistance and stress respons, particularly drought and cold/heat response. We performed GO enrichment on these samples of genes (**Figure S7-S9**), finding enrichment for light stimulus, pollen recognition, and several cellular and metabolic processes. We also assessed whether genes were shared between *Fst* and sweep analyses, hypothesizing that selection on genes within one race could contribute to genetic divergence between races. To perform this analysis, we identified genes shared between *Fst* and CLR analyses - e.g., we compared the set of 396 genes from Mexican-Lowland *Fst* analysis to the 436 CLR genes in the Lowland sample. We found 10 genes shared between the two lists, which was an enrichment relative to random expectation (permutation *p* < 0.0028). The number of shared genes between *Fst* and CLR analyses were higher than expected for four of six comparisons (**Table S5**). The shared genes again constitute another set of credible domestication genes (**Table S8**).

### Contrasting A and B flowering types

To date, the genetic causes of heterodichogamy have not been identified in any system (Endress, 2020). For that reason, we thought it worthwhile to explore genetic factors associated with type A and B flowers. Many of our resequenced accessions had known flowering types, with samples of *n*=13 A types and *n*=9 B types in total (**Table 1**). Importantly, within each flowering type, the samples traversed genetic groups - e.g., the A types included samples from each of the three races (Mexican, Guatemalan and Lowland), and the B types were distributed across Mexican, Guatemalan and hybrid samples. Given the distribution of A and B types across groups, we thought that contrasting the two samples may provide preliminary insights into genomic regions that contribute to this interesting phenotype, perhaps without being overly confounded by population structure.

We therefore performed *Fst* analyses between the two groups, producing a plot with peaks of differentiation between types (**Figure 5A**). The average value of *Fst* was low (at 0.038) compared to *Fst* differentiation between races (average *Fst*: Guatemalan vs. Mexican = 0.241, Guatemalan vs Lowland = 0.223; Lowland vs. Mexican = 0.325) (**Figure S6**), reflecting again the fact that the A vs. B samples do not represent highly differentiated samples. There were nonetheless regions of visually compelling *Fst* peaks between flowering types - e.g., evident peaks on chromosomes 6 and 10, among others. These peaks could be an artifact of the population histories of the samples, but they may also contain genes that differentiate the A and B morph. Consistent with the latter interpretation, the set of 466 genes within the top 1% of *Fst* windows (**Table S9**) were enriched for functions -- like pollination, floral development and photperiodism -- that likely contribute to heterodichogamy (**Figure S10**). Given these functional enrichments, we explored the list of 466 genes to find genes related to floral development and timing, yielding several genes with homologs that affect floral development, circadian rhythm, photoperiodism and the production of volatiles (**Table 2**).

**Figure 5:**
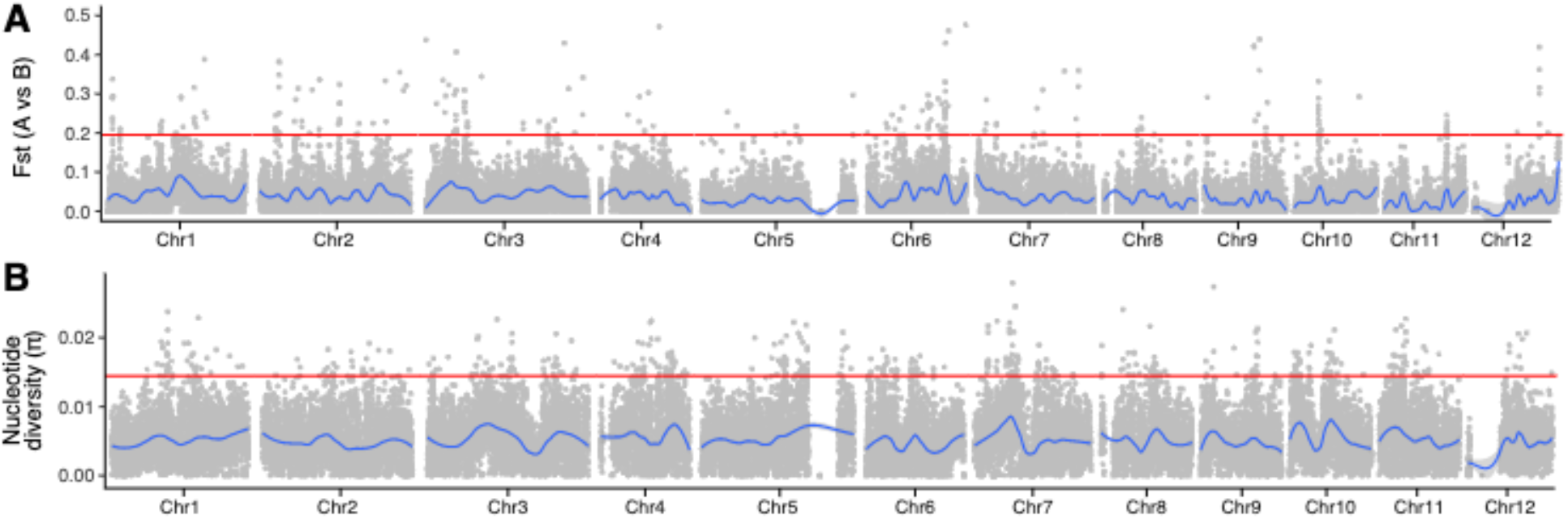
(**a**) A genomic plot of *Fst* windows contrasting the A and B samples of flowering types. The top 1%*Fst* peaks include 466 genes. (**b**) A plot of nucleotide diversity across the 12 pseudo-chromosomes based the ‘pure’ Guatemalan (n = 10) sample.

**Table 2:**
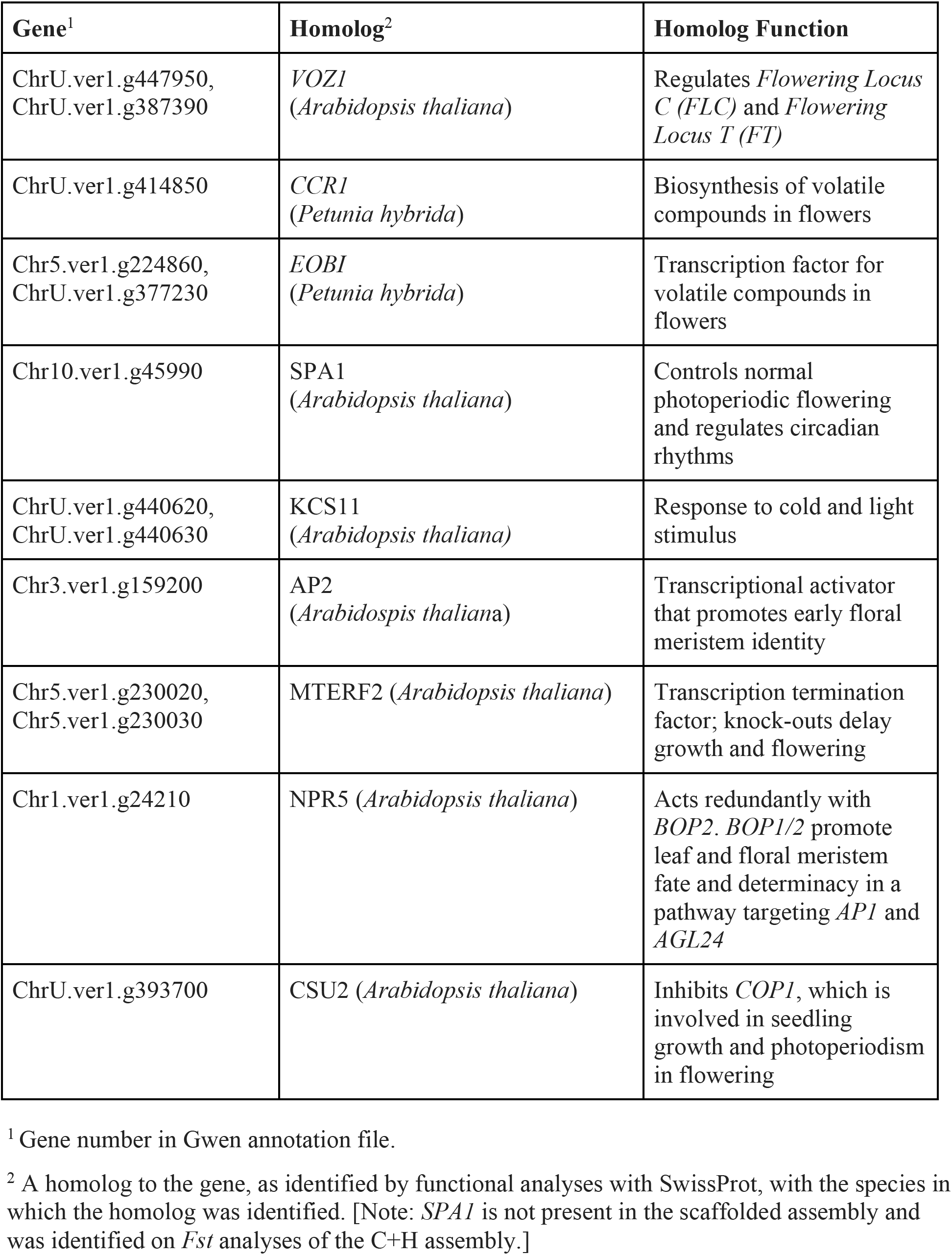
Candidate genes found in *Fst* peaks between samples of A- and B-type flowering accessions with apparent functions in flower development.

We sought another piece of evidence to provide additional support for any of the genes in Table 2. We first wondered whether heterodichogamy could be caused by dosage effects and therefore investigated whether any of the 466 *Fst*-based genes were hemizygous in Gwen, an A-type flower morph. Of the 466 genes, 17 were hemizygous, a percentage (3.6%) nearly identical to the genome average (3.8%), suggesting hemizygosity was not a refining criterion. None of the hemizygous genes were related to floral function (**Table 2**). Second, we hypothesized that the A vs. B polymorphism has been subject to balancing selection, given that heterodichogamy predates the diversification of *P. americana* (Renner, 2001). If true, we expected causal genomic region(s) to have especially high levels of nucleotide diversity within a sample that contained both flowering types. We examined the Guatemalan sample to scan for 5kb regions of high nucleotide diversity. While there were some weakly apparent peaks of diversity, their locations did not correspond with *Fst* peaks between A and B flowering types (**Figure 5B**); none of the genes in Table 2 were among the set of 1% most diverse genes. Overall, three genes were found in both *Fst* peaks and high diversity windows -- a Leucine Rich Repeat gene (Geneid Chr6.ver1.g275850) and two genes similar to the mitochondrial transcription termination factor *MTEF2* (Geneids Chr5.ver1.g230020 and Chr5.ver1.g230030).

## DISCUSSION

We have assembled and annotated the genome of Gwen avocado, which is more complete and contiguous than previously published avocado genomes (Rendón-Anaya *et al*., 2019; Sharma *et al*., 2021). Annotation identified ~65% of the genome as repetitive with 49,450 genes (**Figure 1B**). The latter number is almost two-fold higher than those predicted on the Hass genome (Rendón-Anaya *et al*., 2019) but approaches the 63,000 genes predicted in a transcriptome analysis (Chabikwa *et al*., 2020). Among genes, 3.8% have been detected as hemizygous, due to structural variants that affect > 20% or more of the coding region. This genic hemizygosity value is intermediate among a range that includes perennial grapevines (>12%) (Zhou *et al*., 2019; Vondras *et al*., 2019), an outcrossing annual relative of rice (8.9%) and selfing rice (<<1%) (Kou *et al*., 2020). Given that avocado is clonally propagated, we naively expected hemizygosity to be similar to grapevine. The low value for Gwen probably reflects low divergence between its parents and perhaps the fact that the sampled Gwen tree does not yet have an extensive history of clonality, which may promote maintenance of hemizygous deletions. We do know that genic hemizygosity contributes to trait differences among grapevines (Carbonell-Bejerano *et al*., 2017); it will be interesting to monitor similar issues in avocado for accessions like Mendez, a somatic variant of Hass.

We generated the Gwen genome as a tool for breeding but used it here for investigating questions about the evolutionary history of *P. americana* – e.g., as a reference to address questions about the number of domestication events and the timing of divergence among botanical races. Our analyses are consistent with three independent domestication events (Furnier *et al*., 1990; Ashworth & Clegg, 2003; Rendón-Anaya *et al*., 2019), because PCA, admixture and phylogenetic analyses clearly differentiated among the three botanical races (**Figures 2 & 3**). We also estimated divergence times (**Figure 3**), which varied from ~40,000 years between Lowland and Guatemalan races to > 1.0 My between Mexican and the two other races. These estimates are much older than the expected domestication times for perennial crops (Miller & Gross, 2011; Gaut *et al*., 2015), and hence likely reflect divergence among wild lineages that eventually became sources for domestication.

Our analyses have also provided insights into the inter-racial hybrid origins of some cultivars, representing the first whole-genome insights for most of our samples. Many of our results confirmed findings based on microsatellite and other marker types (Ashworth & Clegg, 2003; Chen *et al*., 2008, 2009) - e.g., accessions like Zutano and Bacon were previously thought to be hybrids between races, which was confirmed in our analyses (**Table S2**) - but also offered surprises. The most notable was the genetic history of Hass, which was traditionally thought to be of Guatemalan origin (Ashworth & Clegg, 2003) but had been inferred to be roughly 50% Guatemalan and 50% Mexican from genetic analyses (Chen *et al*., 2009; Rendón-Anaya *et al*., 2019). We find Hass falls squarely into the Guatemalan group, as do the close relatives of Hass in our sample **(Figure S2)**. Although we do not know the source of disagreement among studies, the genetic provenance of Hass - the most cultivated accession in the world - may not yet be fully resolved.

We removed hybrid accessions to define “pure” samples for population genetic analyses. The resulting sizes of samples varied among races, and likely contributed to some of the variance in results among races. For example, the small sample size of the Mexican sample (*n*= 3) likely led to false positives for sweep genes (638 CLR genes) compared to the Guatemalan sample (*n*=10 with 92 CLR genes). We have nonetheless found a few compelling patterns. For example, the set of high-CLR genes in the Guatemalan race was enriched for functions related to fruit development, likely a trait targeted for domestication. This gene set is, then, fitting for further study and has the potential to help disentangle the origin of some agronomic traits. In addition, some genes were shared as putative domestication genes more often than expected, including 18 genes between the Lowland and Mexican groups. In theory, these genes could represent genomic regions that are particularly prone to maintaining history of sweeps (i.e., low recombination regions) but genome-wide patterns of the CLR statistic do not suggest this is the cause (**Figure 4**). We consider it more likely that these genes represent parallel selection pressures for independent domestication events, but we have few insights into how they may have contributed to domestication traits.

We used similar approaches to investigate regions of genomic divergence between samples defined either by race or by flowering type. The latter yielded the most promising insights, representing (to our knowledge) the first attempt to define genomic regions that may contribute to heterodichogamy. Our inferences are at best preliminary, but they have yielded some interesting candidates, including homologs to: *VOZ1*, which regulates *FLC*, a transcription factor that functions as a repressor of floral transition and contributes to temperature compensation of the circadian clock (Mitsuda *et al*., 2004; Yasui *et al*., 2012); *SPA1*, which contributes to the regulation of circadian rhythms of flowering processes in *A. thaliana* (Ishikawa *et al*., 2006); and *APETALA2*, which binds to thousands of loci in the developing flower and controls various aspects of floral development and organ identity (Yant *et al*., 2010). Yet, none of these genes bear an obvious signal of balancing selection, which is something we hypothesize should be present for a long-lived genetic polymorphism that contributes to A vs. B flowering types. The few genes that do overlap between diversity and divergence seem unlikely to affect heterodichogamy [although it should be noted that MTEF2 knock-outs affect flowering and plant growth (Lee *et al*., 2021)]. There are several reasons why it might be difficult to detect a balancing polymorphism, if there is indeed a balanced polymorphism, but these candidate genes nonetheless suggest a way forward to study this interesting biological phenomenon. Future work will assess the segregation patterns of polymorphisms within candidate genes in a larger sample of A vs. B avocado accessions and also search for trans-specific polymorphism (Charlesworth, 2006) across species of the Lauraceae, since heterodichogamy is also found in other Lauraceae species (Renner, 2001).

## ACKNOWLEDGEMENTS

R. Gaut generated resequencing libraries, G. Gaut provided analysis insights, J. Ross-Ibarra provided feedback on ideas. E. Solares was supported by the NSF graduate research fellowship program, the University of California President’s Pre-Professoriate Fellowship and a UC Presidential Postdoctoral Fellowship. This work used the Extreme Science and Engineering Discovery Environment (XSEDE), which is supported by National Science Foundation grant number ACI-1548562. Specifically, it used the Bridges system, which is supported by NSF award number ACI-1445606, at the Pittsburgh Supercomputing Center (PSC). The XSEDE grants were MCB190050 and MCB180035 to ES. This work was supported in part by a USDA grant #2020-04640 to M.L. Arpaia

## AUTHOR CONTRIBUTIONS

ES, AM-C, SW, VA and AM performed analyses; RFB and DC optimized DNA extraction and supervised long-read sequencing; EF and MLA aided sampling. ES, AM-C, EF, MLA and BSG designed the project. ES, AM-C and BSG co-wrote the manuscript with feedback from all authors.

## DATA AVAILABILITY

The data for this study was submitted to NCBI under BioProject: PRJNA758103, which contains all raw PacBio and Illumina sequencing data, as well as the scaffolded genome assembly. A gff file describing the genes and the scaffold annotation files are available at Zenodo: https://doi.org/10.5281/zenodo.6392169. Published resequencing data used in this study from Rendon-Anaya et al (2019) were from NCBI numbers: SRR8295599, SRR8295600, SRR8295601, SRR8295602, SRR8295603, SRR8295604, SRR8295605, SRR8295607, SRR8295608, SRR8295609, SRR8295610 and SRR8295611.The RNAseq data used for gene annotation were downloaded from NCBI with accession numbers SRR6116327, SRR6116328, SRR6116329, SRR6116330, SRR2000042. The scripts used for analyses are available from https://github.com/GautLab/avo_ref_paper

