## Supplemental File for "Insights into the domestication of avocado and potential genetic contributors to heterodichogamy"

The following Supporting Information is available for this article:

**Fig. S1** PCAs for all samples used in our analysis.
**Fig. S2** Additional admixture plots based on including all *P. americana* accessions and some subsets.
**Fig. S3** The outcome of GO enrichment analysis for the pure Guatemalan sample (n =10), based on the set of 92 candidate genes (Table S4) detected by selective sweep mapping.
**Fig. S4** The outcome of GO enrichment analysis for the pure Lowland sample (n =5), based on the set of 436 candidate genes (Table S4) detected by selective sweep mapping.
**Fig. S5** The outcome of GO enrichment analysis for the pure Mexican sample (n =3), based on the set of 683 candidate genes (Table S4) detected by selective sweep mapping.
**Fig. S6** Plots illustrating *Fst* across the 12 scaffolded pseudo-chromosomes.
**Fig. S7** The outcome of GO enrichment analysis based on the set of 401 candidate genes (Table S7) detected by *Fst* divergence analysis between the Mexican and Lowland pure sample.
**Fig. S8** The outcome of GO enrichment analysis based on the set of 394 candidate genes (Table S7) detected by *Fst* divergence analysis between the Mexican and Guatemalan pure samples.
**Fig. S9** The outcome of GO enrichment analysis based on the set of 385 candidate genes (Table S7) detected by *Fst* divergence analysis between the Lowland and Guatemalan pure samples.
**Fig. S10** The outcome of GO enrichment analysis based on the set of 466 genes (**Table S9**) detected by *Fst* divergence analysis between the A and B Flowering Types.

**Table S1** Assembly Metrics compared across avocado genome assemblies
**Table S2** A list of accessions used in diversity studies with their historical and current assignment to botanical races.
**Table S3** Number of biallelic SNPs and nucleotide diversity per base pair ($\pi$) within the pure samples of botanical races.
**Table S4** List of genes under CLR peaksin each of the three samples representing botanical races. (see Supplementary Excel File)
**Table S5** The outcome of enrichment analyses for comparing genes either inferred to be under selection between two races by Fst and sweep mapping
**Table S6** Lists of 20 genes under CLR peaks that are shared between races with functional annotation information. (see Supplementary Excel File)
**Table S7** List of genes under Fst peaks in pairwise comparisons (see Supplementary Excel File)
**Table S8** List of the 45 genes that were identified as being under both 1% *Fst* peaks and under

1% CLR peaks, with functional annotation information.
**Table S9** List of 466 genes that were under Fst peaks when comparing A flowering types to B flowering types and that were the basis for GO enrichment analysis. (see Supplementary Excel File)

**Methods S1** A brief description of generating mapping and coverage masks for demographic analyses.

**Fig. S1** Additional PCAs based on SNPs. **Top:** The PCA includes all samples used in our analysis, including outgroups. Located in the far left are the *Persea* outgroups and in the top is the *Octea* outgroup. In green just left of the avocados is *Persea scheideanna,* labelled CH-GU-01. **Bottom:** avocado samples including P. *scheideanna* (CH-GU-01), showing it groups clearly within the avocado samples. Keys for both graphs: M = Mexican, L = Lowland, G=Guatemalan, CR = Costa rican, Psch = *P. scheideanna,* labels with crosses are apparent hybrids, N/A = not available (outgroups)

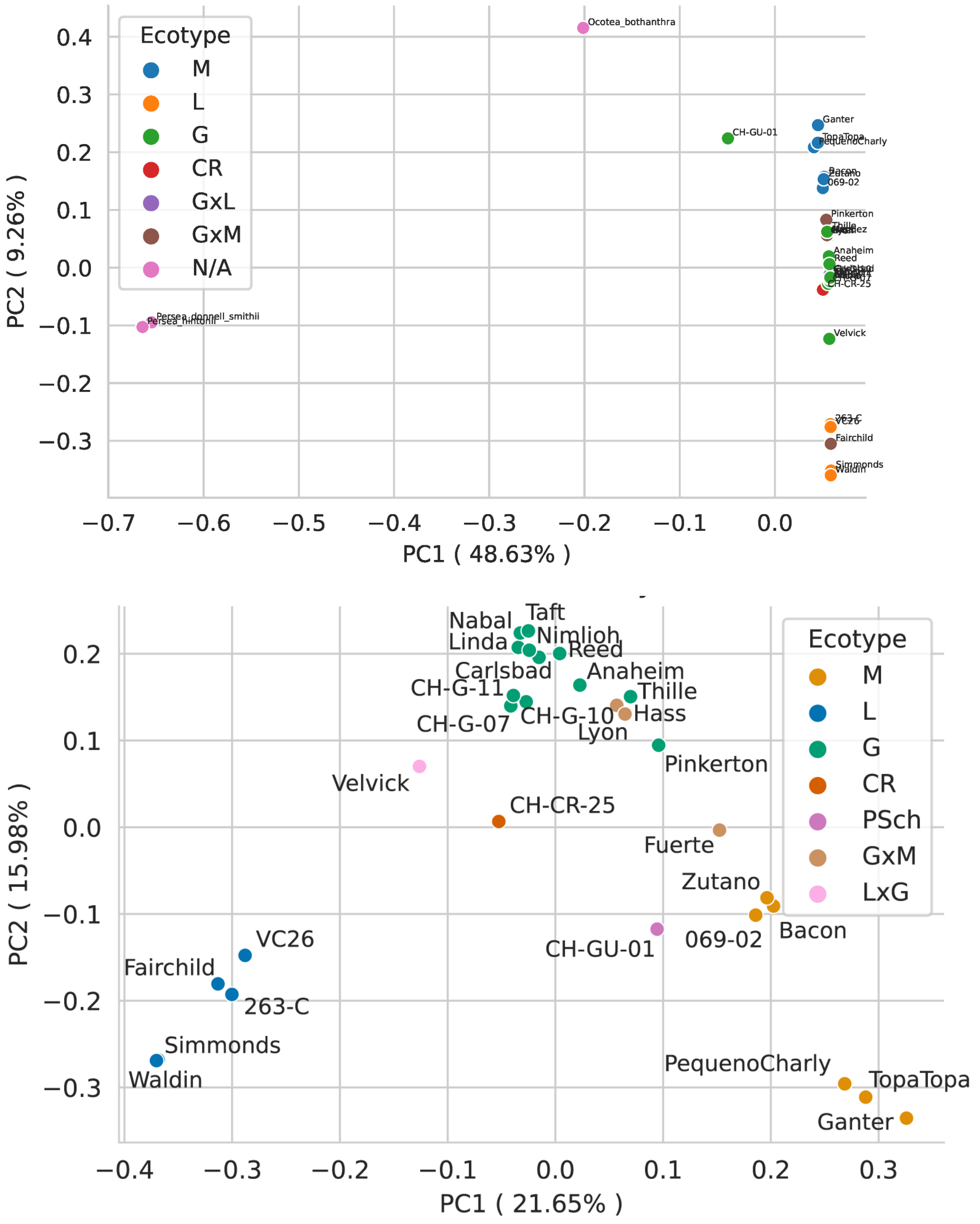

**Fig. S2 Top two graphs:** Admixture plots based on including all *P. americana* accessions, including close relatives of Hass, like Mendez (a purported somatic mutation of Hass), Gwen (a grandchild of Hass), and Thille (an offspring of Hass). With these samples, the optimal grouping is K=4 (middle graph), corresponding to Guatemalan, Mexican, Lowland and Hass groups. The Hass group belongs in the Guatemalan group with K=3 (top graph). **Bottom graph:** Avocado samples with *P. scheideanna* (CH-GU-01) and K=3, reinforcing that it appears to be a hybrid and may not be a good outgroup for evolutionary analyses.

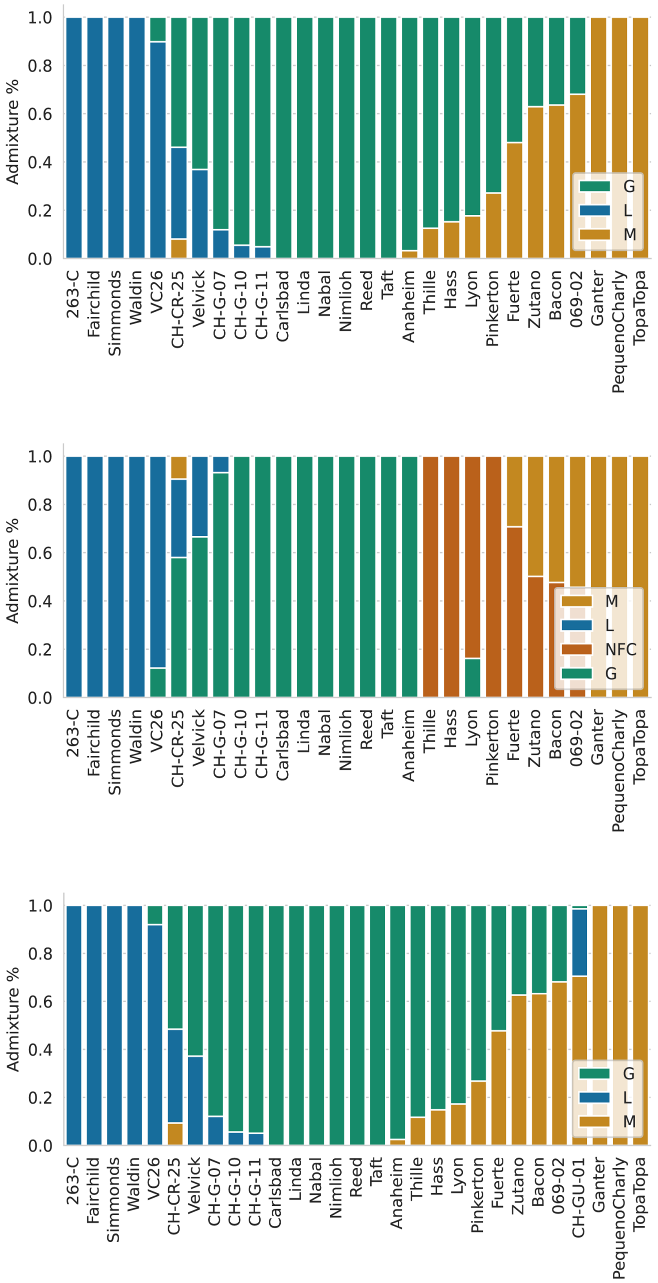

**Fig. S3** The outcome of GO enrichment analysis for the pure Guatemalan sample (n =10), based on the set of 92 candidate genes (Table S4) detected by selective sweep mapping with SweeD. Both graphs were generated by blast2GO, and both include only significant categories as measured by a p-value of $p < 0.05$ by a Fisher's Exact Test, as corrected for multiple tests by the blast2GO program. The graphs differ in the specificity; the top graph was generated to include general terms, while the lower graph used the "reduce to most specific option" to report terms at the most specific level in the gene ontology enrichment directed acyclic graph (DAG) file.

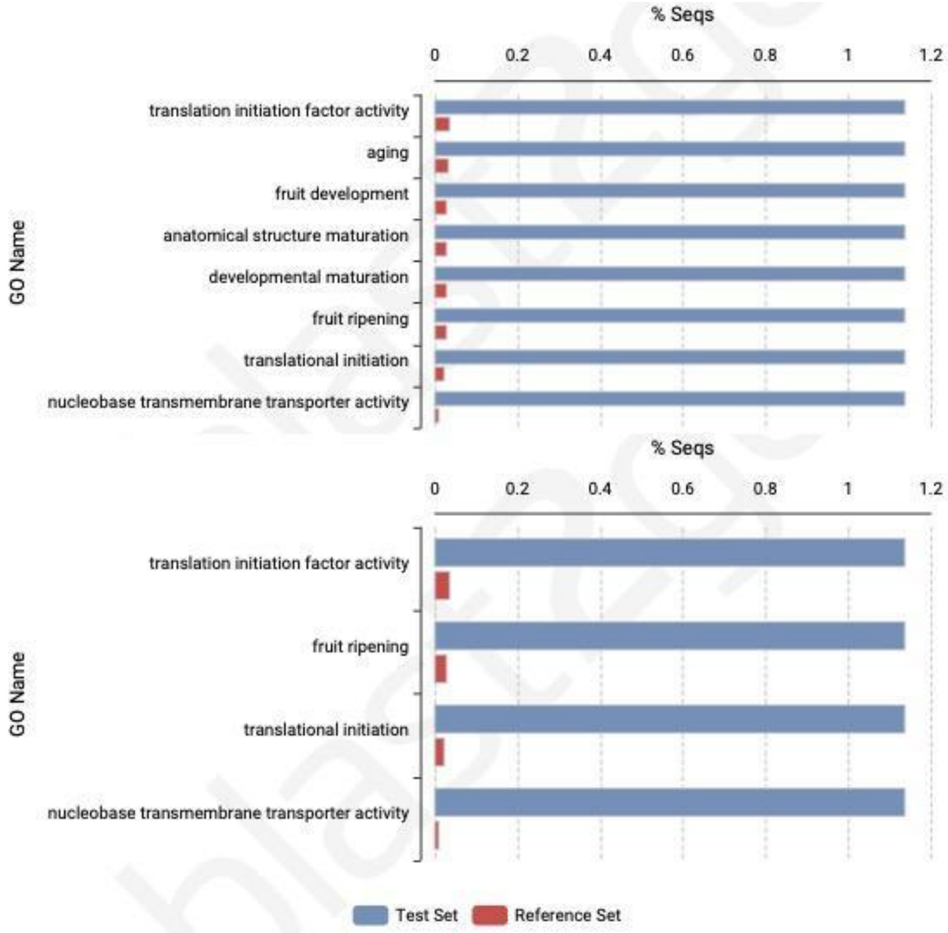

**Fig. S4** The outcome of GO enrichment analysis for the pure Lowland sample (n =5), based on the set of 436 candidate genes (Table S4) detected by selective sweep mapping with SweeD. Both graphs were generated by blast2GO, and both include only significant categories as measure by a p-value of $p < 0.05$ by a Fisher's Exact Test, as corrected for multiple tests by the blast2GO program. The graphs differ in the specificity; the top graph was generated to include general terms, while the lower graph used the "reduce to most specific option" to report terms at the most specific level in the gene ontology enrichment directed acyclic graph (DAG) file.

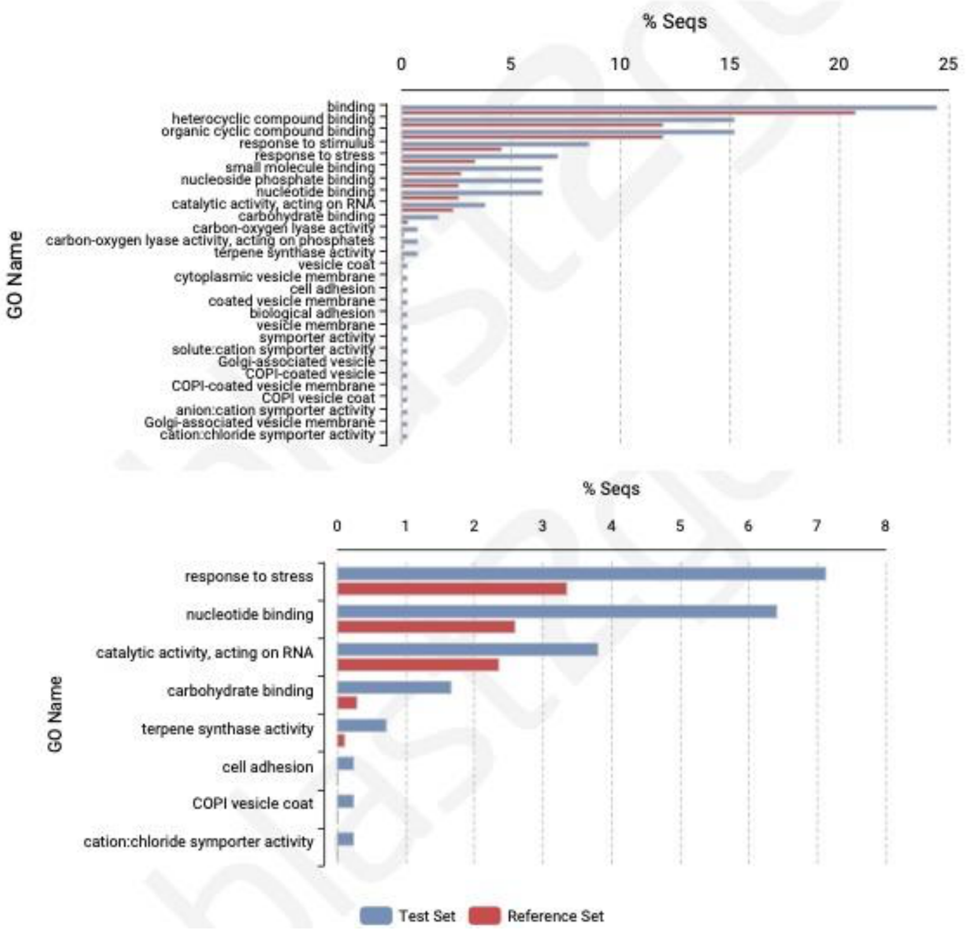

**Fig. S5** The outcome of GO enrichment analysis for the pure Mexican sample (n =3), based on the set of 638 candidate genes (Table S4) detected by selective sweep mapping with SweeD. Both graphs were generated by blast2GO, and both include only significant categories as measure by a p-value of $p < 0.05$ by a Fisher's Exact Test, as corrected for multiple tests by the blast2GO program. The graphs differ in the specificity; the top graph was generated to include general terms, while the lower graph used the "reduce to most specific option" to report terms at the most specific level in the gene ontology enrichment directed acyclic graph (DAG) file.

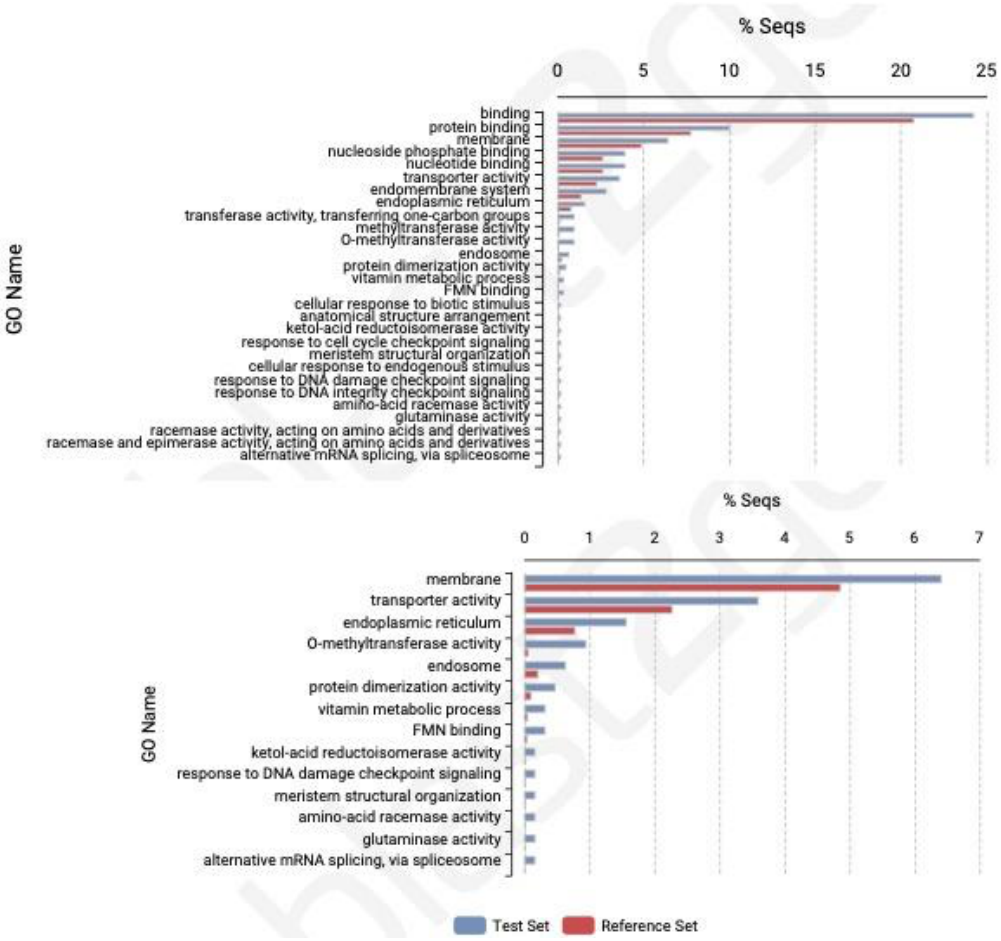

**Fig. S6** Plots illustrating Fst across the 12 scaffolded pseudo-chromosomes. Each plot represents a comparison between two racial samples, based on 20kb windows along the chromosomes. In each graph, a dot represents Fst for each window, the red line represents a smoothed value along the chromosome, and the horizontal blue dotted line indicates the 1% cut-off. The three graphs are as labeled – i.e., the top graph contrasts the Mexican and Lowland sample, the middle graph contrasts the Mexican and Guatemalan sample, and the bottom graph the Lowland and Guatemalan samples.

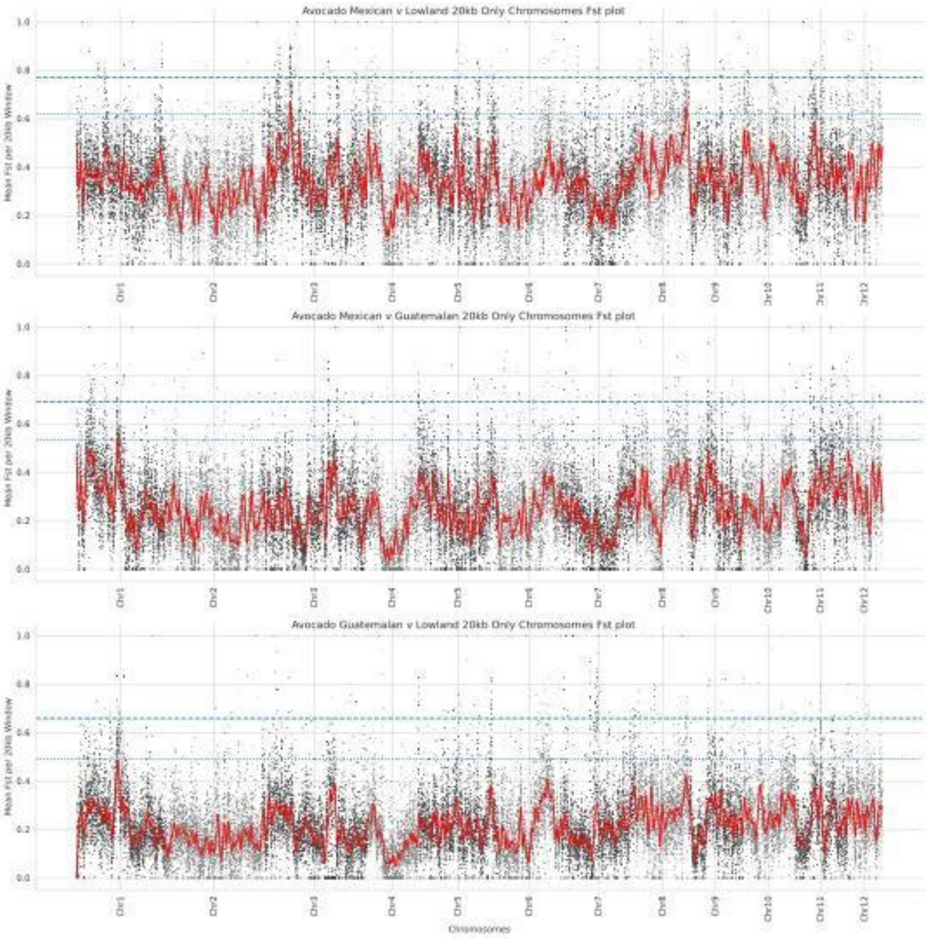

**Fig. S7** The outcome of GO enrichment analysis based on the set of 396 candidate genes (Table S7) detected by Fst divergence analysis between the Mexican and Lowland pure sample. Both graphs were generated by blast2GO, and both include only significant categories as measure by a p-value of $p < 0.05$ by a Fisher's Exact Test, as corrected for multiple tests by the blast2GO program. The graphs differ in the specificity; the top graph was generated to include general terms, while the lower graph used the "reduce to most specific option" to report terms at the most specific level in the gene ontology enrichment directed acyclic graph (DAG) file.

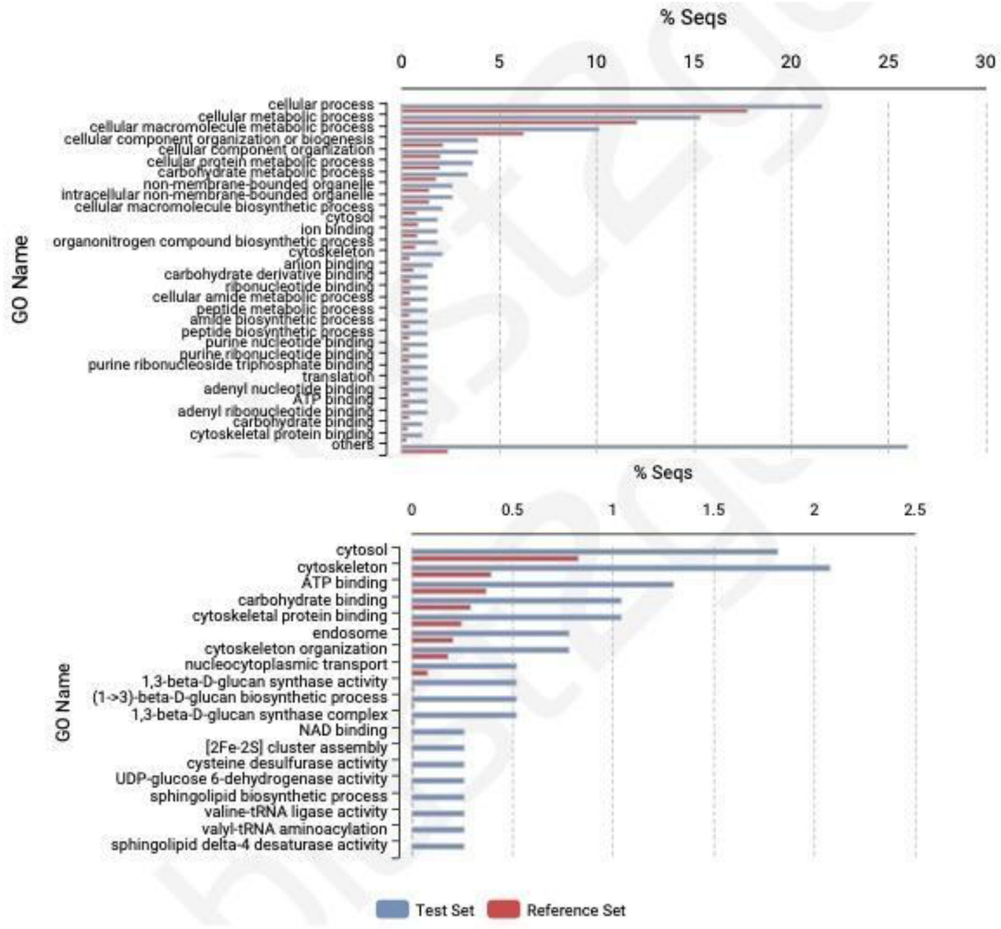

**Fig. S8** The outcome of GO enrichment analysis based on the set of 387 candidate genes (Table S7) detected by Fst divergence analysis between the Mexican and Guatemalan pure samples. Both graphs were generated by blast2GO, and both include only significant categories as measure by a p-value of $p < 0.05$ by a Fisher's Exact Test, as corrected for multiple tests by the blast2GO program. The graphs differ in the specificity; the top graph was generated to include general terms, while the lower graph used the "reduce to most specific option" to report terms at the most specific level in the gene ontology enrichment directed acyclic graph (DAG) file.

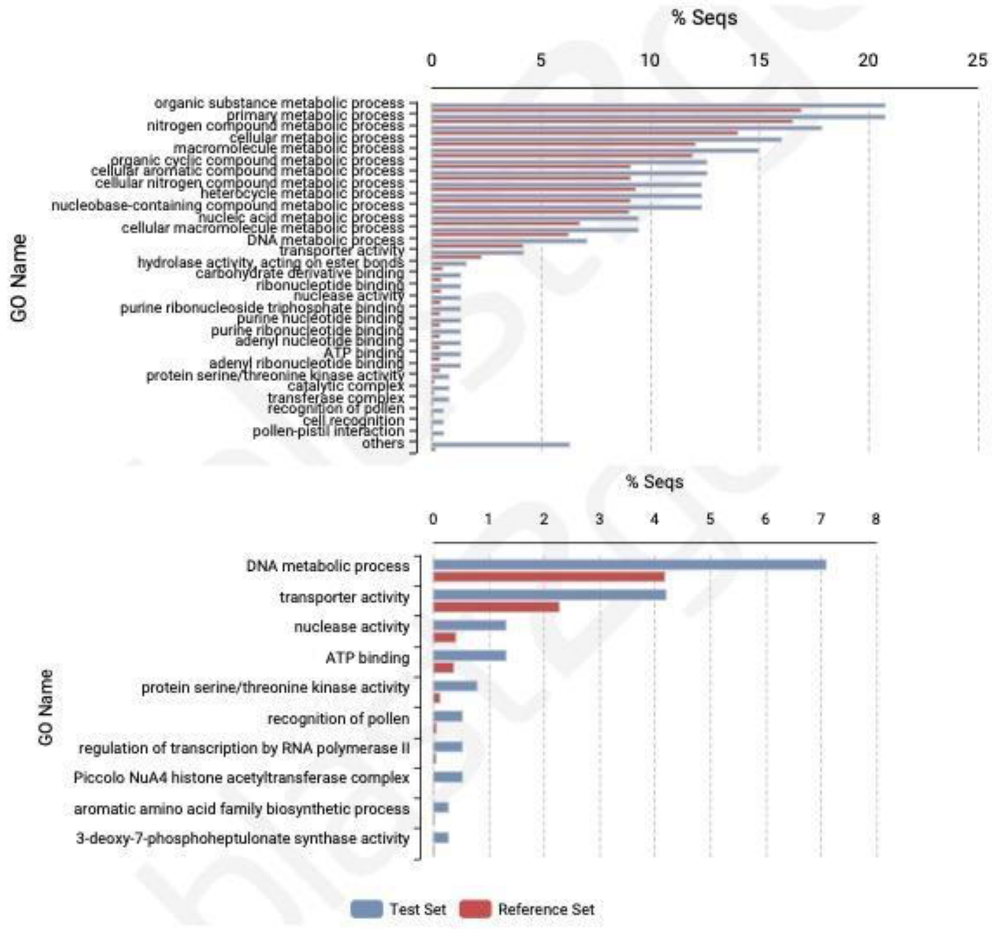

**Fig. S9** The outcome of GO enrichment analysis based on the set of 384 candidate genes (Table S7) detected by *Fst* divergence analysis between the Lowland and Guatemalan pure samples. Both graphs were generated by blast2GO, and both include only significant categories as measured by a p-value of $p < 0.05$ by a Fisher's Exact Test, as corrected for multiple tests by the blast2GO program. The graphs differ in the specificity; the top graph was generated to include general terms, while the lower graph used the "reduce to most specific option" to report terms at the most specific level in the gene ontology enrichment directed acyclic graph (DAG) file.

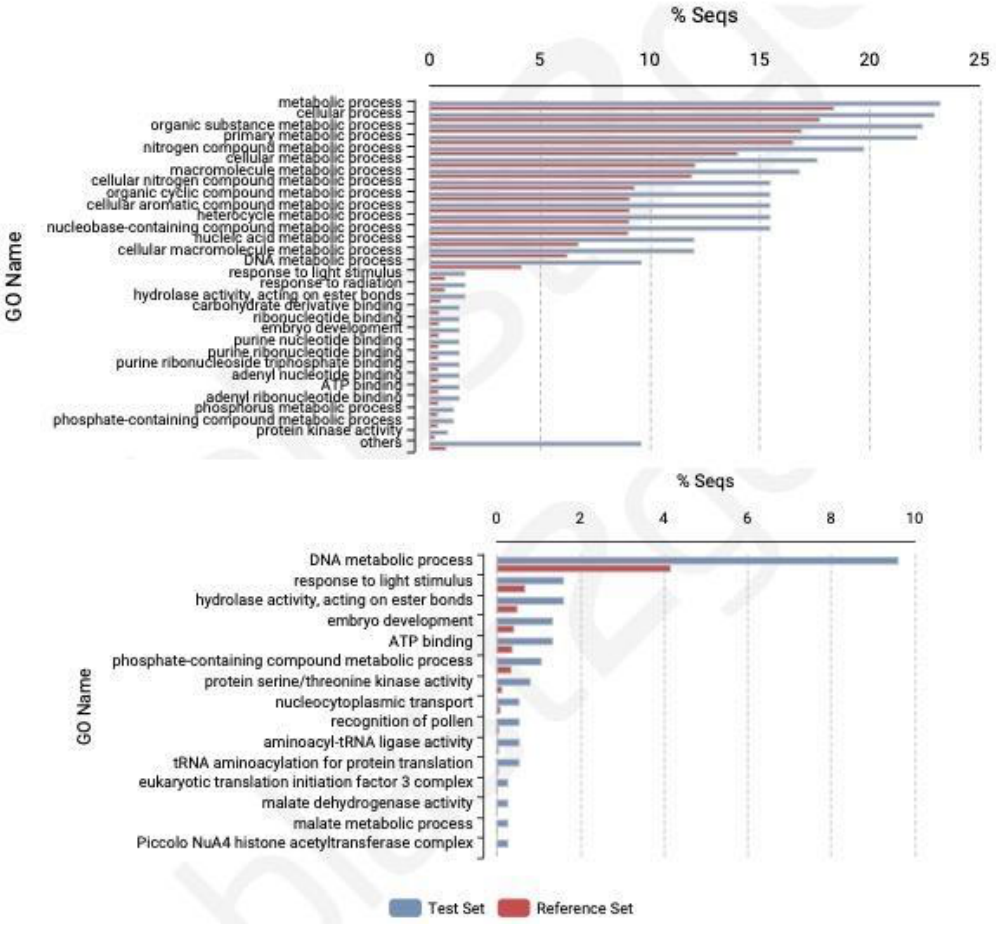

**Fig. S10** The outcome of GO enrichment analysis based on the set of 466 genes (**Table S9**) detected by *Fst* divergence analysis between the A and B flowering types (Table 1). Both graphs were generated by blast2GO, and both include only significant categories as measured by a p-value of $p < 0.05$ by a Fisher's Exact Test, as corrected for multiple tests by the blast2GO program. The graphs differ in the specificity; the top graph was generated to include general terms, while the lower graph used the "reduce to most specific option" to report terms at the most specific level in the gene ontology enrichment directed acylic graph (DAG) file.

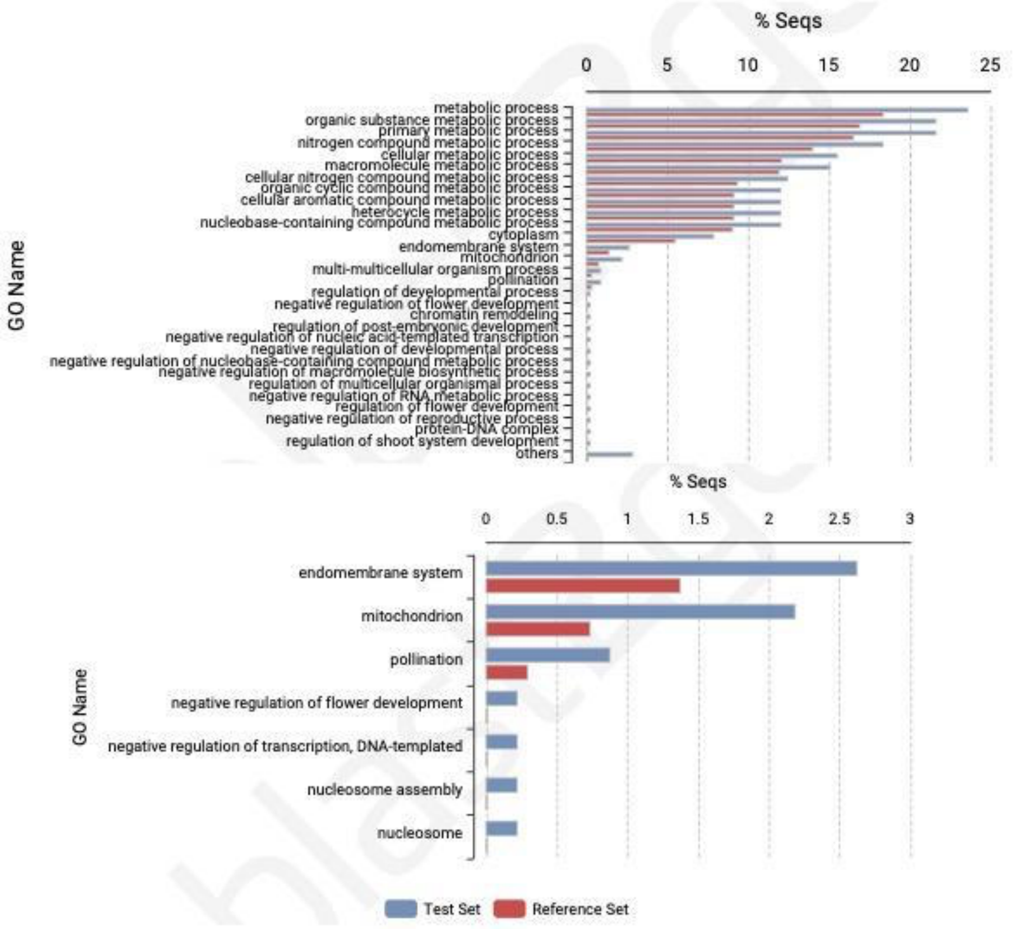

**Table S1** Assembly Metrics for two Gwen assemblies, the Hass HiFi assembly, the Hass assembly and the drymifolia assembly. The Hass HiFi assembly is from (Sharma et al. 2021), and the Hass and drymifolia assemblies are from (Rendón-Anaya et al. 2019)

| | Gwen + HapSolo Contigs | Gwen Scaffolds | Hass HiFi Contigs[1] | Hass Contigs | *drymifolia* Contigs |
|---|---|---|---|---|---|
| **Assembly Size (Mb)** | 1,032 | 703 | 749 | 913 | 823 |
| **Number Fragments** | 989 | 12 | 298 | 8,135 | 42,722 |
| **Largest (kb)** | 17,080 | 85,354 | 12,461 | 2,811 | 4,611 |
| **Percent of Assembly in 12 Largest** | 17.00% | 78.20% | 12.84% | 2.74% | 4.32% |
| **Percent in Fragments > 10Mb** | 17.00% | 78.20% | 5.24% | 0.00% | 0.00% |
| **N50 (Mb)** | 3.37 | 61.89 | 4.33 | 0.30 | 0.32 |
| **L50** | 176 | 5 | 57 | 770 | 502 |
| **BUSCO (%)** | 90.7 | 87.9 | 90.7 | 84.9 | 86.3 |

[1] These values differ from the original publication, based on reanalysis of the publicly available assembled genome, using the same methods we used to evaluate the Gwen genome assemblies.

**Table S2** A complete list of accessions, corresponding to Table 1 in the main text. Historical ecotype assignment follows Rounds (1950), if listed, or later sources if not listed. Alternative assignments are noted, if different. CR = var. costaricensis, G = Guatemalan, L = Lowland, M = Mexican.

| Accession, Variety or Species | Historical Ecotype | PCA Ecotype | Flowering Type | Origin |
|---|---|---|---|---|
| 069-02 | M[1] | GxM | N/A | Unknown |
| 263-C | L[1] | L | N/A | Hunucma, Yucatan, Mexico |
| Anaheim | G | G | A | Anaheim, California, 1910 |
| Bacon | M[2] (GxM[3]) G[4] | GxM | B | Buena Park, California, 1928 |
| Carlsbad | G | G | A | Mexico, budwood collected 1912 |
| CH-CR-25 | CR[1] | GxL | N/A | Matapalo, Puntarenas, Costa Rica |
| CH-G-07 | G[1] | G | N/A | San Cristobal de las Casas, Chiapas, Mexico |
| CH-G-10 | G[1] | G | N/A | Olanca, Chiapas, Mexico |
| CH-G-11 | G[1] | G | N/A | Olanca, Chiapas, Mexico |
| Fairchild | GxL | L | A | Coconut Grove, Florida, seedling of Collin Red, 1925 |
| Fuerte | GxM (M[4]) | GxM | B | Atlixco, Mexico, budwood collected 1911 |
| Ganter | M | M | B | Whittier, California, 1905 |
| Gwen | GxM[2] (G[4]) | G | A | SCREC, Irvine, California, late 1960s |
| Hass | G (GxM[1,2,4]) | G | A | La Habra Heights, California |
| Linda | G | G | B | Antigua, Guatemala, budwood collected 1914 |
| Lyon | G (GxM[2]) (M[4]) | G | B | Hollywood, California, 1908 |
| Mendez | GxM[5] | G | A | La Habra Heights, California, 1926 |
| Nabal | G | G | B | Antigua, Guatemala, budwood collected 1917 |
| Nimlioh | G | G | B | Antigua, Guatemala, budwood collected 1917 |
| *Ocotea botrantha* | N/A | N/A | N/A | Unknown |
| Pequeno Charly | M[1] | M | N/A | Unknown |
| *Persea donnell-smithii* | N/A | N/A | N/A | Unknown |
| *Persea hintonii* | N/A | N/A | N/A | Unknown |

| | | | | |
|---|---|---|---|---|
| *Persea schiedeana* (CH-GU-01) | N/A | GxM | N/A | Mazatenango, Suchitepequez, Guatemala |
| Pinkerton | GxM[2] (G[4]) | G/GxM | A | Saticoy, California, late 1960s |
| Reed | G[2] | G | A | Carlsbad, California, putative Anaheim x Nabal progeny, 1960 |
| Simmonds | L | L | A | Miami, Florida, seedling of Pollock, 1908 |
| Taft | G | G | A | Seed from purchased fruit, likely Mexican origin, 1899 |
| Thille | GxM[2] (G[4]) | G | B | Santa Paula, California, 1946 |
| TopaTopa | M | M | A | Ojai, California, 1907 |
| VC26 | L[6] | L | A | Volcani Institute, Israel |
| Velvick | L[7] | GxL | | University of Queensland, Australia |
| Waldin | L | L | A | Homestead, Florida, 1909 |
| Zutano | M (GxM[4]) | GxM | B | Fallbrook, California, 1926 |

[1] Rendón-Anaya et al. (2019)
[2] UCAVO website (http://ucavo.ucr.edu/avocadovarieties/varietyframe.html)
[3] California Avocado Society (1951)
[4] Chen et al. (2009)
[5] Illsley-Granich et al. (2011)
[6] Ben-Ya'acov and Michelson (1995)
[7] Anderson (2004–2005)

**Table S3** Number of biallelic SNPs and nucleotide diversity per base pair (Pi) within the pure samples of botanical races.

| | Sample Size | Pi | Gwen Scaffold Assembly | 12 pseudo-chromosomes (Scaffolds) |
|---|---|---|---|---|
| Guatemalan | 10 | 0.00446 | 15,822,594 | 11871168 |
| Lowland | 5 | 0.00344 | 12,266,227 | 9,080,921 |
| Mexican | 3 | 0.00370 | 11,792,195 | 8,887,373 |

**Table S4** List of genes under CLR peaks, as inferred from SweeD analysis, in each of the three samples representing botanical races (see supplementary excel file TableS4)

**Table S5** The outcome of enrichment analyses for comparing genes either inferred to be under selection between two races (as inferred by SweeD analyses) or identified to be under selection (SweeD) in one race and contributing to diversity (as measured by Fst) between races. Statistically significant results indicate that more genes are shared between candidate lists than expected at random.

| **Comparison**[1] | **Source List 1**[2] | **Source List 2** | **No. Shared Genes**[3] | **p-value**[4] |
|---|---|---|---|---|
| SweeD v SweeD | Lowland | Mexican | 18 | **p < 0.0001** |
| SweeD v SweeD | Mexican | Guatemalan | 2 | p = 0.3692 |
| SweeD v SweeD | Lowland | Guatemalan | 0 | p = 1 |
| Fst v Sweed | Guatemalan_v_Lowland | Guatemalan | 8 | **p < 0.0001** |
| Fst v Sweed | Guatemalan_v_Lowland | Lowland | 12 | **p = 0.0003** |
| Fst v Sweed | Mexican_v_Guatemalan | Mexican | 5 | p = 0.6354 |
| Fst v Sweed | Mexican_v_Guatemalan | Guatemalan | 6 | **p < 0.0001** |
| Fst v Sweed | Mexican_v_Lowland | Mexican | 5 | p = 0.6513 |
| Fst v Sweed | Mexican_v_Lowland | Lowland | 10 | **p = 0.0028** |

**1** Each comparison contrasts two lists of genes from either two SweeD analysis or from a SweeD analysis and an Fst Analysis.
2 The list of genes from Fst analyses are denoted by, e.g., Mexican_v_Lowland, whereas genes from SweeD analyses are listed by race only.
3 The number of genes shared between lists.
4 The p-value, based on permutation, under the null hypothesis that the number of shared genes is random. Bolded are significant at alpha < 0.05 after multiple test correction.

**Table S6** List of 20 genes under CLR peaks that are shared between races, with information from functional annotation. Only the number shared between Lowland and Mexican samples were higher than random expectation.

| GeneID | Sources[1] | Protein Category[2] | GO activity[3] |
|---|---|---|---|
| Chr1.ver1.g27910 | lowland, mexican | Retrovirus-related Pol polyprotein from transposon TNT 1-94 (EC 3.1.13.-) | hydrolase activity [GO:0016787]; nucleic acid binding [GO:0003676]; zinc ion binding [GO:0008270]; DNA integration [GO:0015074] |
| Chr10.ver1.g47660 | guatemalan, mexican | Uncharacterized protein | |
| Chr10.ver1.g47670 | guatemalan, mexican | | |
| Chr11.ver1.g80970 | lowland, mexican | Hexosyltransferase (EC 2.4.1.-) | glycosyltransferase activity [GO:0016757] |
| Chr12.ver1.g88360 | lowland, mexican | DNA-directed RNA polymerase I subunit rpa49 | nucleolus [GO:0005730]; DNA binding [GO:0003677]; DNA-directed 5'-3' RNA polymerase activity [GO:0003899]; transcription, DNA-templated [GO:0006351] |
| Chr12.ver1.g88370 | lowland, mexican | DNA-directed RNA polymerase I subunit rpa49 | nucleolus [GO:0005730]; DNA binding [GO:0003677]; DNA-directed 5'-3' RNA polymerase activity [GO:0003899]; transcription, DNA-templated [GO:0006351] |
| Chr2.ver1.g121020 | lowland, mexican | Mediator of RNA polymerase II transcription subunit 27 isoform X1 | mediator complex [GO:0016592] |
| Chr2.ver1.g121030 | lowland, mexican | | |
| Chr2.ver1.g131740 | lowland, mexican | Mediator of RNA polymerase II transcription subunit 27 isoform X1 | mediator complex [GO:0016592] |
| Chr3.ver1.g191700 | lowland, mexican | | |
| Chr4.ver1.g203230 | lowland, mexican | | |
| Chr4.ver1.g209590 | lowland, mexican | Sugar transport protein 7 (Hexose transporter 7) | endomembrane system [GO:0012505]; integral component of membrane [GO:0016021]; plasma membrane [GO:0005886]; pollen tube [GO:0090406]; arabinose transmembrane transporter activity [GO:0042900]; symporter activity [GO:0015293]; L-arabinose transmembrane transport [GO:0042882] |
| Chr4.ver1.g209600 | lowland, mexican | Sugar transport protein 7 (Hexose transporter 7) | endomembrane system [GO:0012505]; integral component of membrane [GO:0016021]; plasma membrane |

| | | | |
|---|---|---|---|
| | | | [GO:0005886]; pollen tube [GO:0090406]; arabinose transmembrane transporter activity [GO:0042900]; symporter activity [GO:0015293]; L-arabinose transmembrane transport [GO:0042882] |
| Chr4.ver1.g214170 | lowland, mexican | WAT1-related protein | integral component of membrane [GO:0016021]; transmembrane transporter activity [GO:0022857] |
| Chr6.ver1.g269660 | lowland, mexican | | |
| Chr6.ver1.g269670 | lowland, mexican | | |
| Chr6.ver1.g274670 | lowland, mexican | Uncharacterized protein (Fragment) | ribosome [GO:0005840]; structural constituent of ribosome [GO:0003735]; translation [GO:0006412] |
| Chr6.ver1.g274680 | lowland, mexican | Mitogen-activated protein kinase 10-like protein | ATP binding [GO:0005524]; protein kinase activity [GO:0004672] |
| Chr9.ver1.g365220 | lowland, mexican | RS-norcoclaurine 6-O-methyltransferase-like protein | O-methyltransferase activity [GO:0008171]; protein dimerization activity [GO:0046983]; methylation [GO:0032259] |
| Chr9.ver1.g365230 | lowland, mexican | INO80 complex subunit B-like protein conserved region | Ino80 complex [GO:0031011]; chromatin remodeling [GO:0006338] |

[1] Source refers to the samples that shared the listed gene near the CLR peak.
[2] Functional annotation from Swissprot and/or Uniref , where available.
[3] GO categories based on blast2GO analyses.

**Table S7** List of genes under Fst peaks in pairwise comparisons (see supplementary excel files Table S7)

**Table S8** List of the 45 genes that were identified as being under both 1% Fst peaks and under 1% CLR peaks, with functional annotation information.

| GeneID | Source | Protein name2 |
|---|---|---|
| Chr10.ver1.g43710 | MvL_Low | Uncharacterized protein |
| Chr10.ver1.g47470 | GvL_Guat | Uncharacterized protein |
| Chr10.ver1.g47480 | GvL_Guat | Reverse transcriptase Ty1/copia-type domain-containing protein |
| Chr10.ver1.g47490 | GvL_Guat | Reverse transcriptase Ty1/copia-type domain-containing protein |
| Chr11.ver1.g74150 | MvL_Low | Serine/threonine-protein phosphatase 6 regulatory ankyrin repeat subunit B-like protein |
| Chr12.ver1.g85940 | MvG_Guat | Ribonuclease HI (RNase HI) (Sto-RNase HI) (EC 3.1.26.4) |
| Chr12.ver1.g85950 | MvG_Guat | Uncharacterized protein |
| Chr12.ver1.g85960 | MvG_Guat | Uncharacterized protein |
| Chr12.ver1.g86000 | MvG_Guat | Transposon Tf2-6 polyprotein (Retrotransposable element Tf2 155 kDa protein) |
| Chr12.ver1.g86010 | MvG_Guat | |
| Chr12.ver1.g86020 | MvG_Guat | Retrovirus-related Pol polyprotein from transposon opus [Includes: Protease (EC 3.4.23.-); Reverse transcriptase (EC 2.7.7.49); Endonuclease] |
| Chr12.ver1.g88570 | GvL_Low | ABC transporter |
| Chr2.ver1.g131740 | GvL_Low | Soluble starch synthase 3 chloroplastic/amyloplastic-like protein |
| Chr3.ver1.g164320 | MvL_Mex | Putative serine/threonine protein kinase IREH1 isoform X1 |
| Chr3.ver1.g166260 | MvL_Low | |
| Chr3.ver1.g166280 | MvL_Low | |
| Chr3.ver1.g179740 | MvL_Low | |
| Chr3.ver1.g179750 | MvL_Low | Transposon Tf2-6 polyprotein (Retrotransposable element Tf2 155 kDa protein) |
| Chr3.ver1.g193760 | MvL_Mex | F-box/FBD/LRR-repeat-like protein isoform X1 |
| Chr4.ver1.g212850 | MvL_Low | Glutamate receptor 2.5-like protein isoform X1 |
| Chr5.ver1.g216160 | GvL_Low | Tetratricopeptide repeat-containing domain-containing protein |
| Chr5.ver1.g216190 | GvL_Low | Sodium-dependent phosphate transport protein 1, chloroplastic-like protein |
| Chr5.ver1.g236370 | MvG_Mex | Tonoplast dicarboxylate transporter (AttDT) (Sodium-dicarboxylate cotransporter-like) (AtSDAT) (Vacuolar malate transporter) |
| Chr5.ver1.g236380 | MvG_Mex | Putative F-box protein PP2-B12 |
| Chr5.ver1.g256990 | MvL_Low | Uridine monophosphate kinase (EC 2.7.4.22) (Uridylate kinase) |
| Chr6.ver1.g268430 | GvL_Low | |
| Chr6.ver1.g272410 | GvL_Low | RNA-binding protein 1-like protein |
| Chr6.ver1.g275850 | MvG_Mex | Putative leucine-rich repeat receptor-like serine/threonine-protein kinase |

| | | |
|---|---|---|
| Chr6.ver1.g275860 | MvG_Mex | Non-specific serine/threonine protein kinase (EC 2.7.11.1) |
| Chr6.ver1.g276640 | MvL_Mex | DYW_deaminase domain-containing protein |
| Chr7.ver1.g287380 | GvL_Low | Pentatricopeptide repeat-containing protein, mitochondrial |
| Chr7.ver1.g287390 | GvL_Low | Geraniol synthase, chloroplastic |
| Chr7.ver1.g301160 | GvL_Guat | Putative NADP-dependent oxidoreductase domain-containing protein |
| Chr7.ver1.g301170 | GvL_Guat | Serine/threonine-protein kinase D6PK-like protein |
| Chr7.ver1.g301180 | GvL_Guat | Serine/threonine-protein kinase D6PK-like protein |
| Chr7.ver1.g301190 | GvL_Guat | Protein kinase domain-containing protein |
| Chr7.ver1.g301200 | GvL_Guat | Serine/threonine-protein kinase CTR1 (EC 2.7.11.1) (Protein CONSTITUTIVE TRIPLE RESPONSE1) |
| Chr8.ver1.g330460 | MvG_Mex | WD40 repeat |
| Chr8.ver1.g344340 | MvL_Mex | Lipopolysaccharide core biosynthesis mannosyltransferase lpsB |
| Chr8.ver1.g344350 | MvL_Mex | Uncharacterized protein |
| Chr9.ver1.g356100 | GvL_Low | Retrovirus-related Pol polyprotein from transposon RE1 (Retro element 1) (AtRE1) [Includes: Protease RE1 (EC 3.4.23.-); Reverse transcriptase RE1 (EC 2.7.7.49); Endonuclease RE1] |
| Chr9.ver1.g356700 | GvL_Low | Auxin-induced protein |
| Chr9.ver1.g356710 | GvL_Low | Auxin-responsive protein SAUR50 (Protein SMALL AUXIN UP RNA 50) |
| ChrU.ver1.g471560 | GvL_Low, MvL_Low | Putative LRR receptor-like serine/threonine-protein kinase |
| ChrU.ver1.g471570 | MvL_Low | actin-related protein 2/3 complex subunit 1A-like |

[1] Source refers to the comparisons used to identify candidate genes. The first three letters refer to the Fst comparson, and the last phrase refers to the Sweed analysis. E.g., MvL_Low is based on Fst genes found between Mexican (M) and Lowland (L) races and the comparison of those genes to the candidate gene set from Lowland (Low) CLR analyses.
[2] Functional annotation from Swissprot and/or UniRef, where available.

**Table S9** List of 466 genes that were under Fst peaks when comparing A flowering types to B flowering types and that were the basis for GO enrichment analysis (Figure S10) (see supplementary excel file Table S9).

**Methods S1** For demographic analyses, the mappability mask was made with (http://lh3lh3.users.sourceforge.net/snpable.shtml) by generating 150 bp mers in 1 bp increments across the genome, then mapping sequences back to the genome with BWA v0.7.8-r455 (Li and Durbin 2010) and identifying "mappable" regions where the majority of sequences mapped uniquely without mismatches. To include only regions with sufficient sequencing coverage, we calculated the coverage from the alignment file of each sample with the bedcov program from samtools (v1.10) package; we then created a mask per sample that only included regions with sequencing coverage higher than 5x..
